# Bi-specific CAR-iNKT cell immunotherapy for high-risk KMT2A-rearranged leukemia outperforms CAR-T in an NKG2D-dependent manner and eradicates leptomeningeal disease

**DOI:** 10.1101/2025.01.22.633541

**Authors:** Hongwei Ren, Natalina Elliott, Bryan Lye, Mohammad Umer Sharif Shohan, Joe W Cross, Lucy Field, Kanagaraju Ponnusamy, Siobhan Rice, Thomas Jackson, Ilia Leontari, Nouhad El Ouazzani, Rebecca Thomas, Sarah Inglott, Jack Bartram, Owen Smith, Jonathan Bond, Irene AG Roberts, Christina Halsey, Rachael Bashford-Rogers, Thomas A Milne, Anindita Roy, Anastasios Karadimitris

## Abstract

Current therapies, including autologous CAR-T immunotherapy, fail to cure half of infants with *KMT2A*-rearranged acute lymphoblastic leukemia (KMT2Ar-ALL). Here we deploy allogeneic iNKT cells, ‘innately’ more powerful effectors than T cells, and equip them with CD19- and/or CD133-targeting CARs. Compared to mono-specific counterparts and bi-specific CAR-T, CD19-CD133 bi-specific CAR-iNKT have more potent anti-leukemia activity, they effectively target CAR antigen-low leukemia, eradicate medullary and leptomeningeal leukemia and induce sustained remissions without discernible hematologic toxicity. Mechanistically, dynamic CAR- and CAR antigen-dependent upregulation of the activating innate receptor NKG2D and its engagement by corresponding ligands on KMT2Ar-ALL cells lead to more potent anti-leukemia effect of CAR-iNKT over CAR-T cells, including against CAR antigen-negative leukemia. Thus, by engaging with two different types of leukemia-associated targets, CAR-iNKT provide a powerful platform for the treatment of KMT2Ar-ALL. This approach can be readily adapted for other high-risk malignancies, including those with otherwise difficult to target leptomeningeal involvement.

---

Acute leukemias initiated by rearrangement and fusion of the *KMT2A* (*MLL*) gene to a variety of partners are amongst the worst prognosis hematological malignancies. *KMT2A*-rearranged (KMT2Ar) B cell acute lymphoblastic leukemia (B-ALL) is the commonest form of infant ALL (80%)^1^ and comprises a small fraction of childhood (3%) and adult (10%) ALLs as well^2,3^. Compared to the >90% survival of childhood B-ALL, event free survival of infants and children with KMT2Ar B-ALL, is 38% and 65% respectively, mainly due to chemo-resistance and relapse^4,5^. Recent use of bi-specific CD19-targeting T cell engagers appears to improve survival while CD19 CAR-T cell immunotherapy offers promise of rescuing a subset of patients with relapsed disease who would otherwise have a dismal prognosis^6,7^. Nevertheless, after CD19 CAR-T therapy, one third of patients relapse with either CD19+ or CD19-disease, including lineage-switched disease^8,9^. The latter describes a state of transcriptional plasticity where trans-differentiation or expansion of pre-existing KMT2Ar myeloid/multipotent progenitors, results in B lineage lymphoblasts assuming a myeloid phenotype associated with partial or complete loss of CD19 expression^8,10^ and escape from CD19 CAR-T control^8,11^. Treatment failure in KMT2Ar-ALL is also linked to inability of current therapeutic approaches to effectively eradicate central nervous system (CNS) disease, usually manifesting as leptomeningeal infiltration^12^.

We previously showed that *PROM1* (CD133) is a direct target of leukemic KMT2A fusion proteins and that KMT2Ar-ALL typically express PROM1/CD133^13,14^. Therefore, dual targeting of CD19 and CD133 provides a rational approach to enhance anti-leukemic activity and limit immune escape. In support of this, a pre-clinical approach involving a tandem CD19-CD133 CAR has been described^15^; however, direct comparison of the bi-specific approach with CD133 CAR-T and *in vivo* study of potential hematological toxicity were not performed. This is particularly important as CD133 is also expressed on normal hematopoietic stem and progenitor cells (HSPC)^16-18^. Of note, previous pre-clinical data suggest that tandem CARs may be less effective than bi-specific designs^19,20^, possibly due to steric hindrance mechanisms that interfere with high affinity binding of each CAR to their target antigens.

As well as optimal CAR design and target selection, effective immunotherapy would also benefit from the co-operative activity of powerful and multi-functional effector cells. In this regard, we and others have developed the CD1d-restricted, glycolipid-reactive iNKT cells^21-23^ as a versatile ‘off-the-shelf’ platform^24-27^ that not only lacks the risk of inciting acute graft versus host disease (aGVHD)^28,29^, but also has inherent anti-tumor activity that may complement that of the CAR modules. Notably, we have previously shown that when iNKT are equipped with CAR, they are able to eradicate brain lymphoma more effectively than their CAR-T counterparts^26,30^.

Here, we describe the development of iNKT cells equipped with a bi-specific CAR against CD19 and CD133, and investigate their activity against high risk medullary and meningeal KMT2Ar-ALL.

## Results

### Enhanced anti-leukemic activity of bi-specific CAR-iNKT

Immunophenotypic analysis of patient KMT2Ar ALL primary blast cells, in agreement with our previous study^14^, showed co-expression of CD19 and CD133, with a 67% median level of CD133 co-expression (range 0-100%; **Suppl Figure 1a&b).** Although it was previously reported that KMT2Ar-ALL primary blasts express CD1d^31^, we found that the majority of the patient samples we tested were negative for CD1d expression **(Suppl Figure 1a&c)**. Similarly, ALL blasts isolated from the bone marrow of our recently described KMT2Ar infant ALL model where expression of the *KMT2A::AFF1* fusion gene in human fetal liver CD34+ cells was achieved by CRISPR-Cas9 mediated gene editing (^CRISPR^KMT2A-AFF1 ALL)^32^, demonstrated variable expression of CD133 and minimal expression of CD1d on CD19^+^ blasts **(Suppl Figure 1a&b)**. In contrast, CD133 but not CD19 is expressed on normal primitive fetal hematopoietic stem cell/multipotent progenitors (Lin^-^CD34+CD38^-^) while fetal B-progenitors (CD34^+^CD19^+^) express CD19 but not CD133 confirming that CD133/CD19 co-expression is leukemia-specific **(Suppl Figure 1d).**

Starting from two mAb clones we generated mono-specific 2^nd^ generation 28ζ CD19 and 4-1BBζ CD133 CARs and the corresponding bi-specific CAR such that each CAR would be expressed stoichiometrically and independently of each other in the same cell (**Figure 1a)**. Using our previously described protocol^26,27^, we generated peripheral blood-derived CAR iNKT cells with all three CARs stably and similarly highly expressed. The corresponding CAR-iNKT cells exhibited similar growth rates (**Figure 1b and Suppl Fig 2a)**. When tested against the KMT2A::AFF1 CD19^+^CD133^+^ B-ALL cell line SEM, bi-specific CAR-iNKT were more cytotoxic than their mono-specific counterparts in 4 and 24hr cytotoxicity assays (**Figure 1c).** To test the specificity of each CAR, we generated SEM cell lines that lack expression of either CD19, CD133 or both using CRISPR-Cas9 gene editing (**Figure 1d)**. We tested cytotoxic activity of mono-specific CD19 or CD133 and bi-specific CD19-CD133 CAR-iNKT cells against these and the parental CD133^+^CD19^+^ SEM cells **(Figure 1e)**. As expected, mono- and bi-specific CAR-iNKT displayed the lowest cytotoxicity against CD19^-^CD133^-^ SEM cells while mono-specific CAR-iNKT displayed only background cytotoxicity against SEM cells lacking expression of their respective target. By contrast, bi-specific CAR-iNKT cells displayed the highest cytotoxic activity against the parental CD19^+^CD133^+^ SEM cells, but were also equally effective as the corresponding mono-specific CAR-iNKT cells against CD19^+^CD133^-^ and CD19^-^ CD133^+^ SEM cells **(Figure 1e)**. This effect was also observed in another CD19^+^CD133^+^ KMT2Ar-ALL cell line, RS4;11, where bi-specific CAR-iNKT displayed greater activity compared to their mono-specific CAR counterparts **(Suppl Figure 2b).**

**Figure 1.**
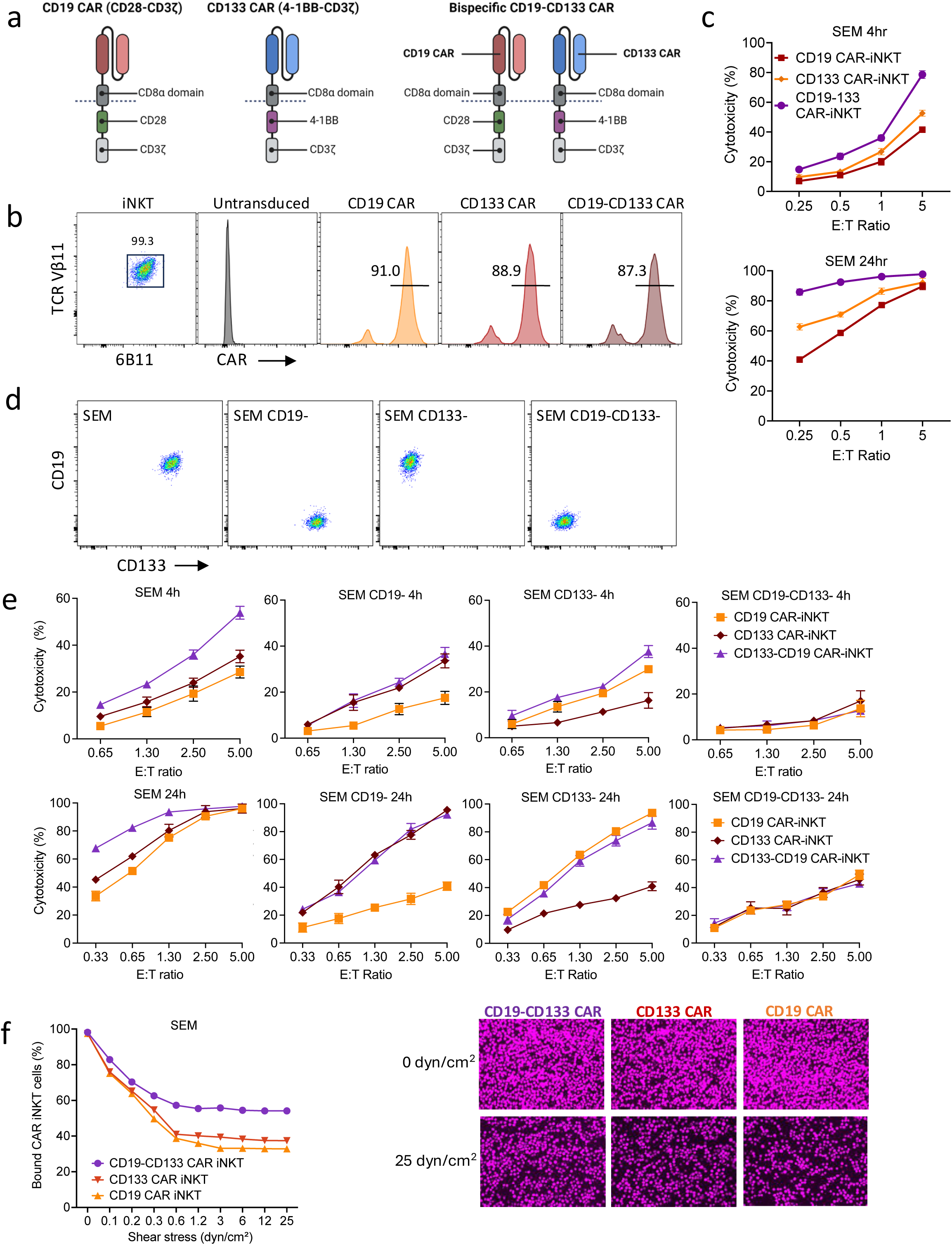
Design and *in vitro* validation of bi-specific CD19-CD133 CAR-iNKT cells. **a.** Design of CD19 and CD133 mono-specific and CD19-CD133 bi-specific CARs. **b.** Flow-cytometric analysis of highly pure, expanded iNKT cells, gated on CD3+ and co-expressing invariant TCRVα24 (identified with the 6B11 mAb) and TCRVβ11, expressing the indicated CARs, as assessed by L-protein staining. **c.** Cytotoxicity assay of indicated mono- and bi-specific CAR-iNKT against SEM cells at 4 and 24 hrs. Mean and standard deviation of triplicate assays is shown. **d.** Flow-cytometric analysis of CD19 and CD133 expression in parental and gene-edited SEM leukemia cells. **e.** Cytotoxicity assay of indicated mono- and bi-specific CAR-iNKT against the target cells shown in d. **f.** Flow chamber avidity assay of indicated CAR-iNKT over a range of pressures (left) and representative images of CAR-iNKT bound to SEM cells at the beginning and end of the assay (right). Data are from two independent experiments using two different iNKT donors.

These findings confirm that both CD19 and CD133 CARs are active as bi- as well as mono-specific CARs when transduced into iNKT cells, with bi-specific CAR-iNKT exerting a more robust anti-leukemia activity than their mono-specific counterparts *in vitro*. This likely reflects increased avidity mediated by their ability to engage both CD19 and CD133 on leukemia cells. To investigate this, we applied a modified flow chamber assay in which avidity was assessed by measuring the fraction of immune cells attached to leukemia cells under an increasing pressure gradient. We found that while mono-specific CD133 CAR-iNKT demonstrated slightly higher avidity than the CD19 counterparts, bi-specific CAR-iNKT displayed the highest avidity **(Figure 1f**).

### Bi-specific CAR-iNKT outperform mono-specific CAR-iNKT *in vivo* and eradicate medullary and meningeal leukemia

We next tested the anti-leukemic efficacy of bi-specific CAR-iNKT cells *in vivo*. First, to confirm the anti-leukemic efficacy of CD133 CAR-iNKT cells, mice were transplanted with luciferase-dsRed-expressing KMT2Ar SEM cells. In this model, mice develop an aggressive form of universally fatal ALL with medullary and extramedullary (spleen, liver and leptomeningeal) disease by day 23-25^33^. We found that CD133 CAR iNKT cells at a dose of 5×10^6^/mouse very effectively controlled leukemia burden as assessed by reduction of bioluminescence activity to background levels (**Suppl Figure 2c)**.

Next, we defined the lowest dose of bi-specific CAR-iNKT cells that could improve survival in mice engrafted with 10^6^ luciferase-dsRed-labelled SEM leukemia cells. We used an SEM subline which after *in vivo* propagation displayed a spectrum of CD19 and CD133 co-expression, from high, low, to no expression of both markers **(Suppl Figure 2d)**. Of the three doses tested, (10^5^, 3×10^5^ and 10^6^ cells), only the highest dose significantly delayed leukemia growth as assessed by bioluminescence imaging (BLI) and significantly prolonged survival **(Suppl Figure 2e&f**). Importantly all the residual leukemia cells while expressing dsRed, were CD19^-^CD133^-^ (**Suppl Figure 2g)** suggesting complete elimination of leukemia cells co-expressing CD19 and CD133 even when expressed at low levels and in line with the higher avidity of bi-specific CAR-iNKT cells.

To test the ability of the bi-specific CAR-iNKT to mitigate immune evasion, we injected NSG mice with 10^6^ parental SEM cells co-expressing CD19 and CD133 admixed with CD19^-^CD133^+^ and CD19^+^CD133^-^ SEM cells at an 8:1:1 ratio respectively, followed by treatment with mono- or bi-specific CAR-iNKT cells at the limiting dose of 10^6^ cells per mouse **(Suppl Figure 3a)**. Treatment with mono-specific as well as bi-specific CAR-iNKT cells resulted in a significant improvement in survival, with bi-specific CAR-iNKT cells conferring the longest survival **(Suppl Figure 3b)**. We performed immunophenotypic analysis of the bone marrow, spleen and liver at termination expecting preferential expansion of the CD19^-^CD133^+^ and CD19^+^CD133^-^ fraction of leukemia cells in animals treated with mono-specific CD19 and CD133 CAR-iNKT respectively but not in those treated with bi-specific CAR-iNKT. Against expectation, with the exception of a significantly higher fraction of CD19^-^CD133^+^ cells in the spleens of animals treated with CD19 CAR-iNKT, we found no significant differences in the frequency of the three different leukemic populations in mice treated with any of the three effectors **(Suppl Figure 3c)**. This suggests a CAR-independent anti-leukemia effect of iNKT cells, shared by all three CAR-iNKT effectors.

Repeating the same experiment using a different iNKT donor and only parental CD19^+^CD133^+^ SEM cells as targets confirmed the enhanced ability of bi-specific CAR-iNKT cells to control leukemia and prolong overall survival when compared to each of the mono-specific CAR-iNKT cells **(Figure 2a&b)**.

**Figure 2.**
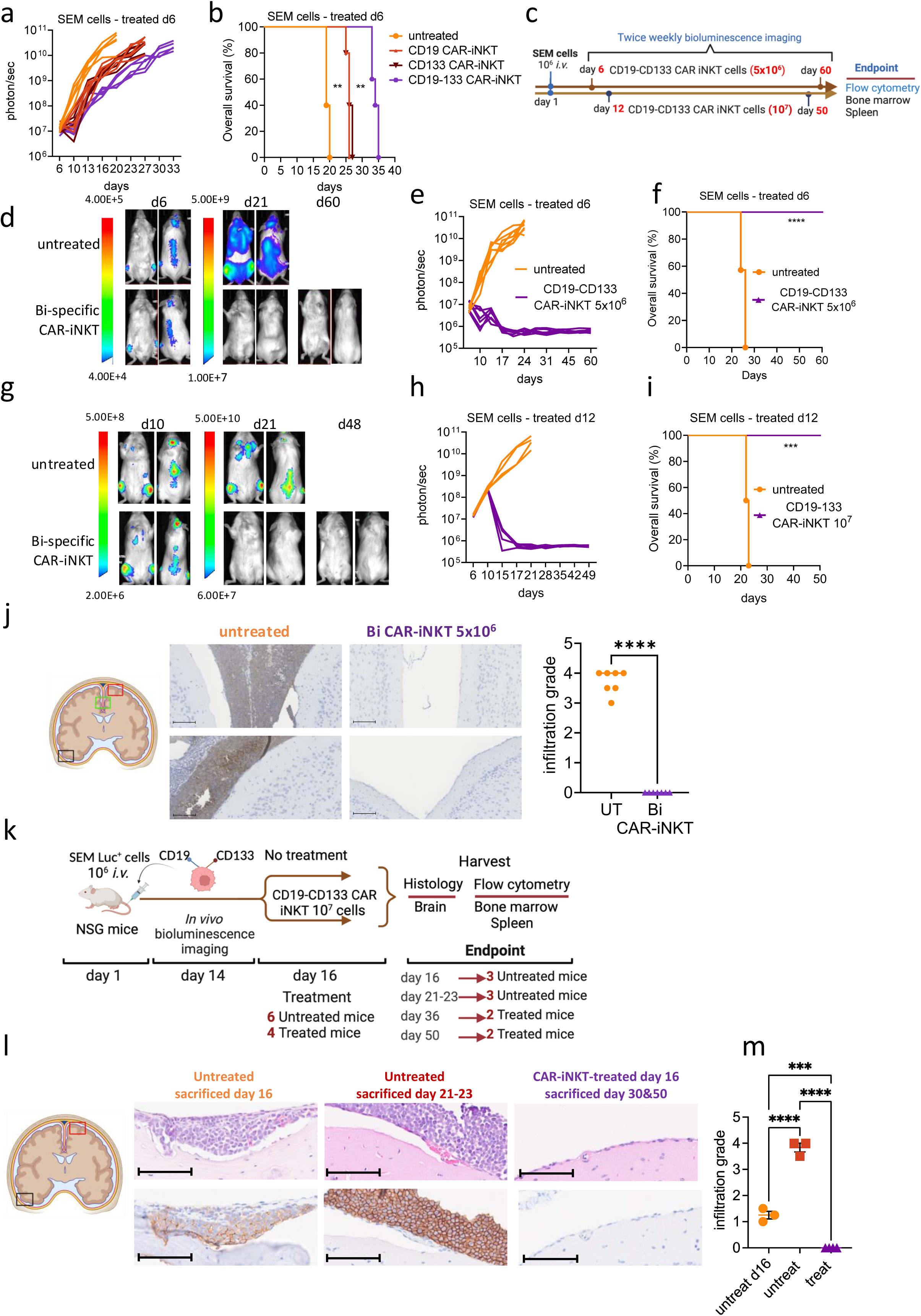
Eradication of medullary and meningeal KMT2Ar ALL by bi-specific CAR-iNKT. **a & b.** Leukemia burden as assessed by bioluminescence imaging (BLI) and overall survival of SEM leukemia-bearing mice treated with 10^6^ mono- or bi-specific CAR-iNKT od day 6 post-leukemia transfer. Kaplan Meier curves were plotted for overall survival with differences assessed by log-rank test (n=5 mice per group). **c.** Schematic of experiments shown in d-f & g-i. **d-f.** Representative BLI images, leukemia burden and overall survival in mice treated with 5×10^6^ bi-specific CAR-iNKT 6 days after iv leukemia cell transfer (n=7 mice per group). **g-i.** Representative BLI images, leukemia burden and overall survival in mice treated with 10^7^ bi-specific CAR-iNKT 12 days after iv leukemia cell transfer (n=5 mice per group). **j.** Coronal sections of representative mouse brains and immunohistochemistry from end of experiment untreated control and end of bi-specific CAR-iNKT 5×10^6^ treated mice (as shown in d-f) stained with anti-human CD19. Imaging region indicated by green box. Scale bar = 100µM. **k**. Left: Schematic of experiment. **l.** Coronal sections of representative mouse brains from untreated mice sacrificed on day 16, untreated mice sacrificed at termination on days 21-23 and from day 16-treated mice sacrificed either on day 36 or 50. Top panel: H&E staining; bottom panel: immunohistochemistry against human CD19. Day 16 area indicated by black box and the other two groups by red box (left). Scale bar = 100µM. **m.** Cumulative meningeal infiltration grade scores in untreated mice sacrificed on day 16, untreated mice sacrificed at termination on days 21-23 and in day 16-treated mice sacrificed either on day 36 or 50. Mann-Whitney or one-way ANOVA. **p<0.01, ***p<0.001, ****p<0.0001

To investigate whether bi-specific CAR-iNKT can truly eradicate established leukemia we treated SEM leukemia-bearing mice with 5×10^6^ (**Figure 2c&d-f)** and 10^7^ CAR-iNKT (**Figure 2c&g-i)** on days 6 and 12 respectively when leukemia burden in the BM was at ∼2% and ∼15% **(Suppl Figure 4a)**. In both experiments, leukemia burden as assessed by BLI rapidly declined to baseline with 100% of animals surviving long term (**Figures 2d-f & g-i)**. Furthermore, detailed analysis of the BM and spleen at termination showed lack of leukemia as well as effector cells **(Suppl Figure 4b)**. Critically, while in untreated animals the meninges were heavily infiltrated by leukemia cells as identified by H&E and CD19 staining of brain sections (infiltration grade >3 out of 5), leukemia cells were not identified in the meninges of CD19-CD133 CAR-iNKT-treated animals **(Fig 2j).**

Therefore, CD19-CD133 bi-specific CAR-iNKT cells can eradicate leukemia from the BM and spleen and protect from meningeal disease thus resulting in deep, lasting remissions. We then investigated whether bi-specific CAR-iNKT cells can eradicate established meningeal leukemia using three leukemia-bearing groups **(Figure 2k)**: one that was untreated and when assessed for meningeal disease on day 16 by detailed immunohistochemical analysis was found to have a mean leukemia meningeal infiltration grade of 1.25, while a second group that was also untreated and at end-point (D21-23) had an infiltration grade of 3.8 **(Figure 2l&m)**. By contrast, a third group of animals treated with bi-specific CAR-iNKT on day 16 showed no evidence of leukemia by both BLI and upon immunophenotypic analysis (i.e., HLA-class I expressing cells) of the BM and spleen respectively (**Suppl Figure 4c-e).** Importantly, consistent with meningeal leukemia eradication, histological and immunohistochemical analysis of the brain of these animals either on day 36 or day 50 showed complete absence of any CD19-positive cells (infiltration grade 0; **(Figure 2l&m)**. Of note, some meningeal stromal thickening was also observed, likely representing areas of old leukemic infiltrates cleared by CAR-iNKT treatment **(Suppl Figure 4f)**. This is in line with our previous finding that CAR-iNKT cells can eradicate brain lymphoma^26^ and consistent with ability of CAR-iNKT cells to efficiently enter the cerebrospinal fluid space likely through the choroid plexus blood–cerebrospinal fluid barrier^34^.

We conclude that bi-specific CAR-iNKT cells outperform their mono-specific counterparts and at high doses display curative potential against medullary and extramedullary disease, underpinned by their enhanced, CAR-dependent avidity. Critically, they also display ability to eradicate as well as protect from meningeal leukemia.

### CAR-iNKT cells outperform their CAR-T counterparts

Next, we compared the anti-leukemic activity of bi-specific CD19-CD133 CAR-iNKT cells with their CAR-T cell counterparts manufactured in parallel for 28 and 14 days respectively, using the same donor **(Suppl Figure 5a)**. CAR-iNKT cells were more cytotoxic than CAR-T cells against the parental and gene-edited KMT2Ar SEM cells and the RS4;11 cells **(Figure 3a, Suppl Figure 5b)** and showed higher avidity **(Figure 3b)**. Consistent with their ‘innate’ functional profile, CAR-iNKT cells, upon co-culture with SEM cells for 4 and 24hrs produced similar peak levels of IFNγ and TNFα as CAR-T cells, but at an earlier time point, and a higher cytotoxic potential as indicated by production of higher amounts of PRF, GRZB and IL-2 compared to CAR-T cells (**Figure 3c & Suppl Figure 5c).** Of note, CD19 and CD133 mono-specific CAR-iNKT are also more cytotoxic against KMT2Ar leukemia cell lines than their CAR-T counterparts **(Suppl Figure 5d).** As SEM and RS4;11 cell lines do not express CD1d (**Suppl Figure 5e** and **Suppl Fig 6a**) the higher anti-leukemic activity of CAR-iNKT is not due to CD1d targeting.

**Figure 3.**
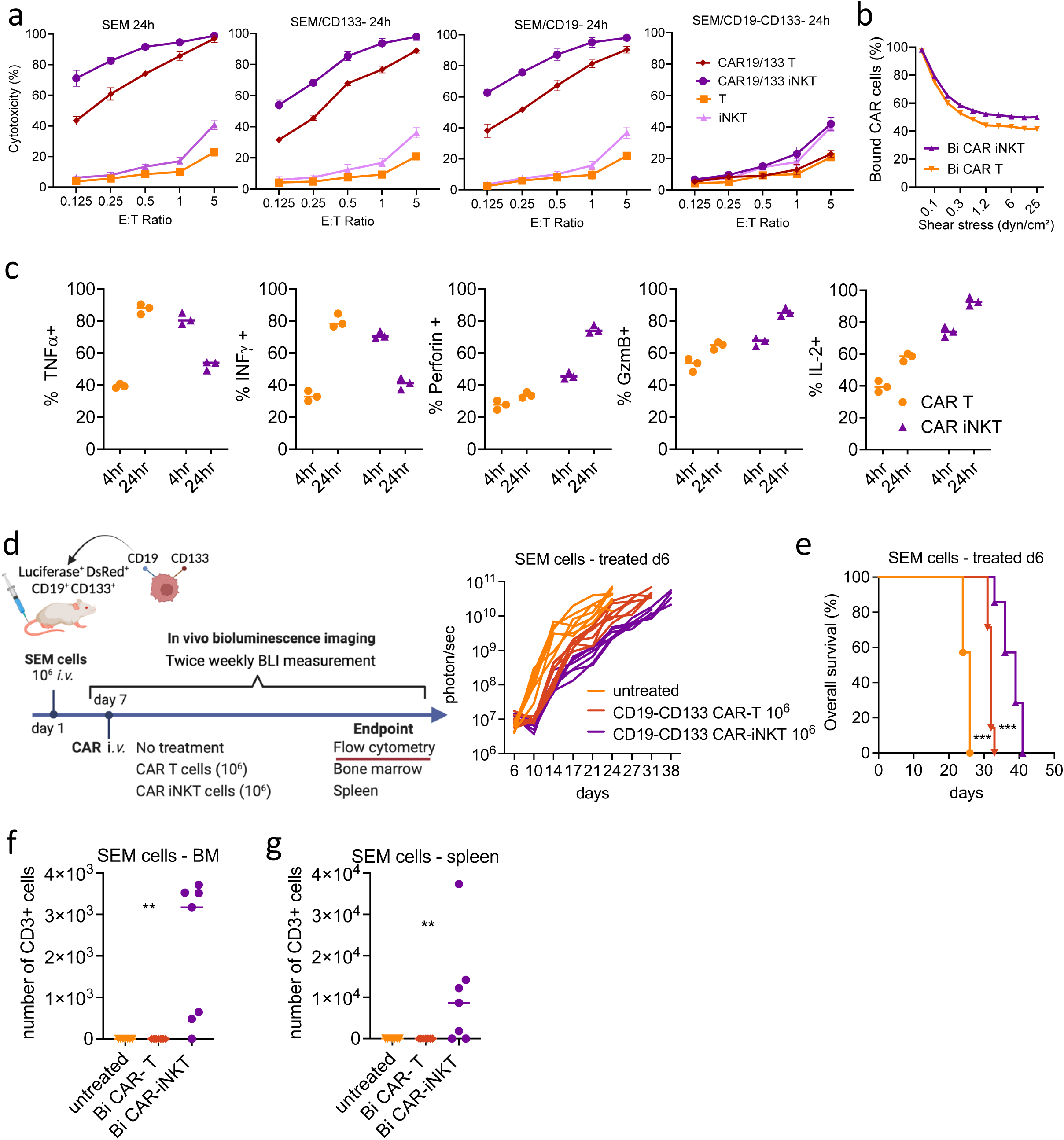
CAR-iNKT outperform CAR-T against KMT2Ar ALL. **a.** Cytotoxic activity at 24hrs of untransduced T and iNKT and of their bi-specific CAR-transduced counterparts against parental SEM cells and their single or double CD19 and CD133 gene edited sublines. **b.** Avidity assay comparing bi-specific CAR-T vs CAR-iNKT against SEM cells. Representative of three independent experiments using three different iNKT donors.. **c.** Expression level of indicated cytokines by bi-specific CAR-T and CAR-iNKT after 4 and 24hr co-culture with SEM leukemia cells. **d&e.** Schematic of experiment**, l**eukemia burden by BLI and overall survival of SEM leukemia-bearing mice treated with indicated dose of bi-specific CAR-T and CAR-iNKT cells (n=7 mice per group). Log rank test, ***p<0.001. **f&g.** Absolute numbers of CAR-T and CAR-iNKT cells in bone marrow and spleen at sacrifice of mice treated as shown in d&e. One-way ANOVA; **p<0.01

Next, we compared the *in vivo* efficacy of bi-specific CAR-T vs CAR-iNKT cells **(Figure 3d)**. We found that at the limiting dose of 10^6^ CAR+ cells, CAR-iNKT-treated SEM leukemia-bearing mice had a significantly lower disease burden and longer survival than CAR-T-treated mice **(Figure 3d&e)**. While CAR-T cells were not detected in the BM and spleen of treated animals at end-point, CAR-iNKT cells were readily detectable at endpoint one week later **(Figure 3f&g)**.

In summary, these findings show that compared to CAR-T, CAR-iNKT cells have higher capacity for *in vivo* persistence and display superior anti-leukemia activity involving a mechanism that is independent of CD1d expression by the leukemia cells.

### Bi-specific CD19-CD133 CAR-iNKT cells eradicate primary KMT2Ar ALL and induce remissions

To better emulate primary KMT2Ar ALL and the pattern of CD19-CD133 co-expression in patient primary blast cells, we took advantage of the ^CRISPR^KMT2A-AFF1 ALL blast cells in which there is negligible CD1d expression **(Suppl Figure 1a&c).** In the ^CRISPR^KMT2A-AFF1 ALL sample used, CD133 was expressed by ∼65% of the CD19+ blasts **(Fig 4a)**.

**Figure 4.**
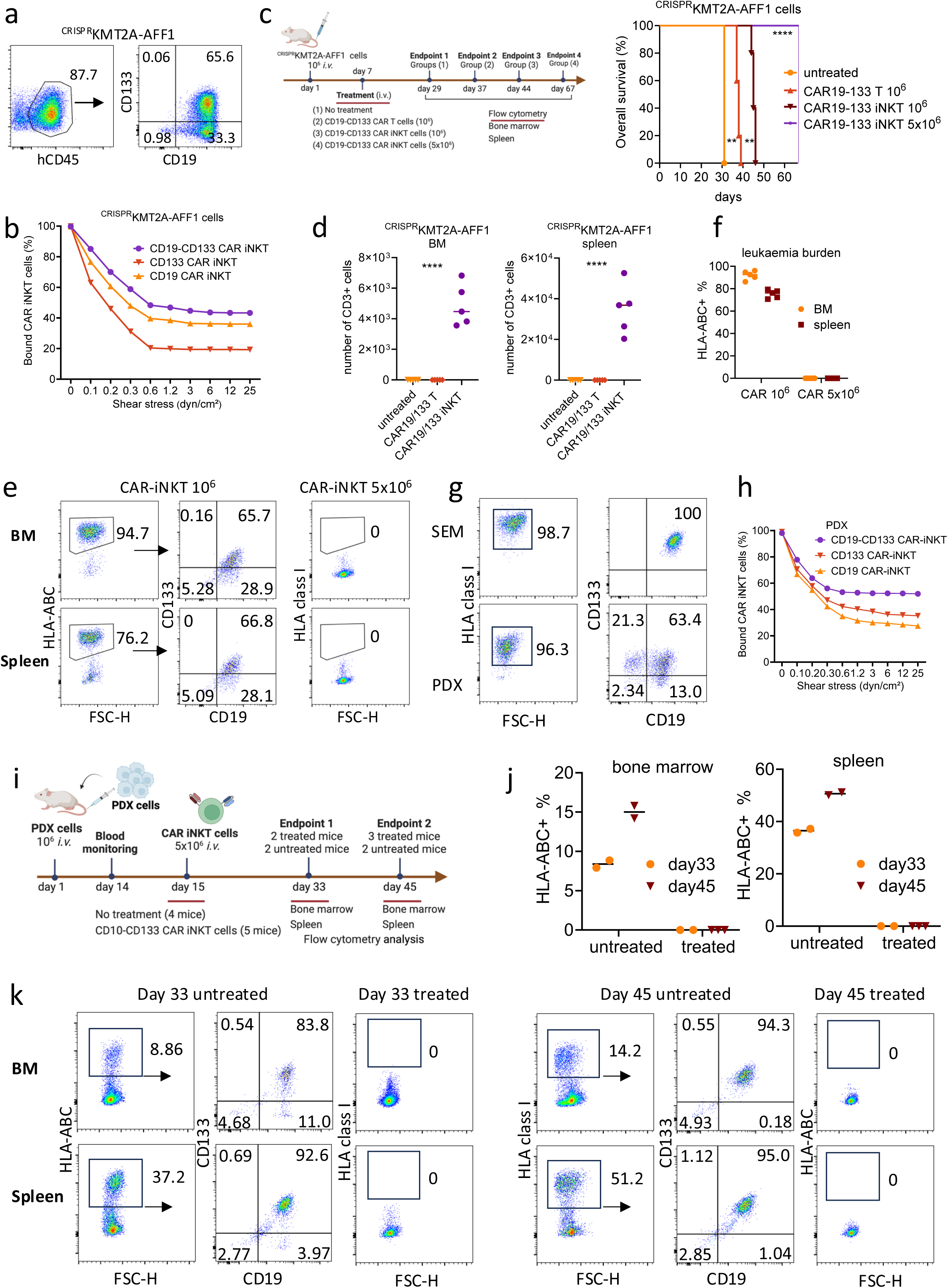
Bi-specific CD19-CD133 CAR-iNKT cells eradicate primary KMT2Ar ALL including of ALL cells lacking CAR target expression. **a.** CD19 and CD133 co-expression pattern in ^CRISPR^KMT2A-AFF1 cells. **b.** Avidity assay of mono- and bi-specific CAR-iNKT against ^CRISPR^KMT2A-AFF1 cells. Data are from two independent experiments and two different iNKT donors.. **c.** Schematic of experiment and overall survival of ^CRISPR^KMT2A-AFF1 leukemia-bearing mice treated with 10^6^ bi-specific CAR-T and CAR-iNKT and 5×10^6^ CAR-iNKT (n=7 mice per group). **d**. iNKT numbers as identified by CD3 expression in BM and spleen of untreated and 10^6^ treated CAR-T and CAR-iNKT. **e.** Flow-cytometric analysis of ^CRISPR^KMT2A-AFF1 leukemia cells as identified with HLA-class I staining in bone marrow and spleen of leukemia-bearing mice treated with 10^6^ or 5×10^6^ bi-specific CAR-iNKT. **f.** Cumulative data of e. **g.** CD19 and CD133 co-expression pattern in PDX KMT2A-AFF1 cells as compared to that of SEM cells. **h.** Avidity assay of indicated CAR-iNKT over a range of pressures. Data are from two independent experiments and two different iNKT donors. **i.** Schematic of experiment and flow-cytometric analysis of leukemia burden in bone marrow and spleen of untreated and 5×10^6^ bi-specific CAR-iNKT-treated PDX KMT2A-AFF1 leukemia-bearing mice 18- and 30-days post treatment. HLA-class I+ cells were gated on 7AAD-live cells**. j.** Cumulative data of i.

As for the KMT2Ar ALL cell lines, bi-specific CAR-iNKT were more cytotoxic against ^CRISPR^KMT2A-AFF1 ALL blasts than CAR-T *in vitro* **(Suppl Figure 5f)** and displayed higher avidity than mono-specific CAR-iNKT **(Figure 4b).**

We treated ^CRISPR^KMT2A-AFF1 leukemia-engrafted mice (assessed by presence of >1% leukemic cells in peripheral blood on day 6; **Suppl Figure 5g)** with 10^6^ bi-specific CAR-iNKT vs CAR-T generated from the same donor **(Figure 4c)**. Similar to SEM cells, we found that ^CRISPR^KMT2A-AFF1 leukemia-bearing mice treated with CAR-iNKT survived significantly longer than CAR-T-treated animals **(Figure 4c)** and while CAR-T were not detectable at sacrifice, CAR-iNKT cells were readily detectable in the BM and spleen of mice sacrificed nearly one week later **(Figure 4d)**. Increasing the therapeutic dose to 5×10^6^ bi-specific CAR-iNKT resulted in 100% of ^CRISPR^KMT2A-AFF1 leukemia-bearing mice surviving beyond 60 days **(Figure 4c).** While untreated and 10^6^ bi-specific CAR-iNKT-treated mice succumbed to disease, in 5×10^6^ bi-specific CAR-iNKT-treated animals there were no detectable HLA-class^+^ cells suggesting leukemia elimination **(Figure 4e&f)**.

Finally, we tested the activity of bi-specific CAR-iNKT against patient-derived xenograft (PDX) leukemia cells derived from an 11-year-old patient with KMT2A-AFF1 ALL. The PDX cells comprised ∼60% CD19^+^CD133^+^ population within the human HLA class I-expressing fraction **(Figure 4g).** *In vitro*, bi-specific CAR-iNKT were more cytotoxic than their CAR-T counterparts **(Suppl Figure 5h)** and demonstrated higher avidity than mono-specific CAR-iNKT **(Figure 4h).**

*In vivo*, animals with detectable PDX cells in blood on day 14 were treated or not with 5×10^6^ bi-specific CAR-iNKT on day 15 **(Figure 4i)**. On day 33, two animals from each arm were sacrificed. While ∼8% and ∼37% HLA class I-expressing cells were detected in the BM and spleen respectively of untreated animals, HLA class I-expressing cells were not detectable in treated animals. Similar analysis of another group of animals sacrificed on day 45 showed that while leukemia burden increased in the bone marrow and spleen of untreated animals, no human cells were detected in treated animals **(Figure 4i&j)**.

We conclude that bi-specific CAR-iNKT persist longer and are more effective than CAR-T against primary ALL, and when delivered in high doses effectively eliminate primary ALL with variable expression of the CAR targets.

### Dynamic NKG2D upregulation in CAR-iNKT cells accounts for their ability to outperform CAR-T cells and target CAR-negative leukemia *in vitro*

We hypothesized that the mechanism that dictates enhanced anti-leukemia activity of CAR-iNKT over CAR-T could involve activating NK receptors such as NKG2D which is known to be expressed on CD4-iNKT cells^35,36^.

To address this hypothesis, we first compared kinetics of NKG2D expression on CAR effectors. We found that in three different donors, expression of NKG2D at baseline was significantly higher (as assessed by both % and expression intensity) in CD19-CD133 CAR-iNKT than CD19-CD133 CAR-T cells (**Figure 5a&b)**. Following a 16hr exposure to SEM or RS4;11 cells, while in CAR-T cells expression of NKG2D increased modestly, in CAR-iNKT cells it increased to nearly 100% (**Figure 5a&b)**. Of note, while as previously reported, only CD4- and not CD4+ iNKT expressed NKG2D at baseline, upon exposure to leukemia cells, both fractions displayed expression of NKG2D to nearly 100%. Next, by analysing previously published transcriptomes of KMT2Ar cell lines (SEM and RS4;11)^37,38^ we found evidence of expression of at least one of the NKG2D ligands (including MICA-B and ULBP1-6) in both cell lines **(Suppl Figure 6a)**. We further investigated expression of NKG2D ligands by staining KMT2Ar cell lines with soluble NKG2D-Fc protein and confirmed expression in all cases tested (**Figure 5c)**.

**Figure 5.**
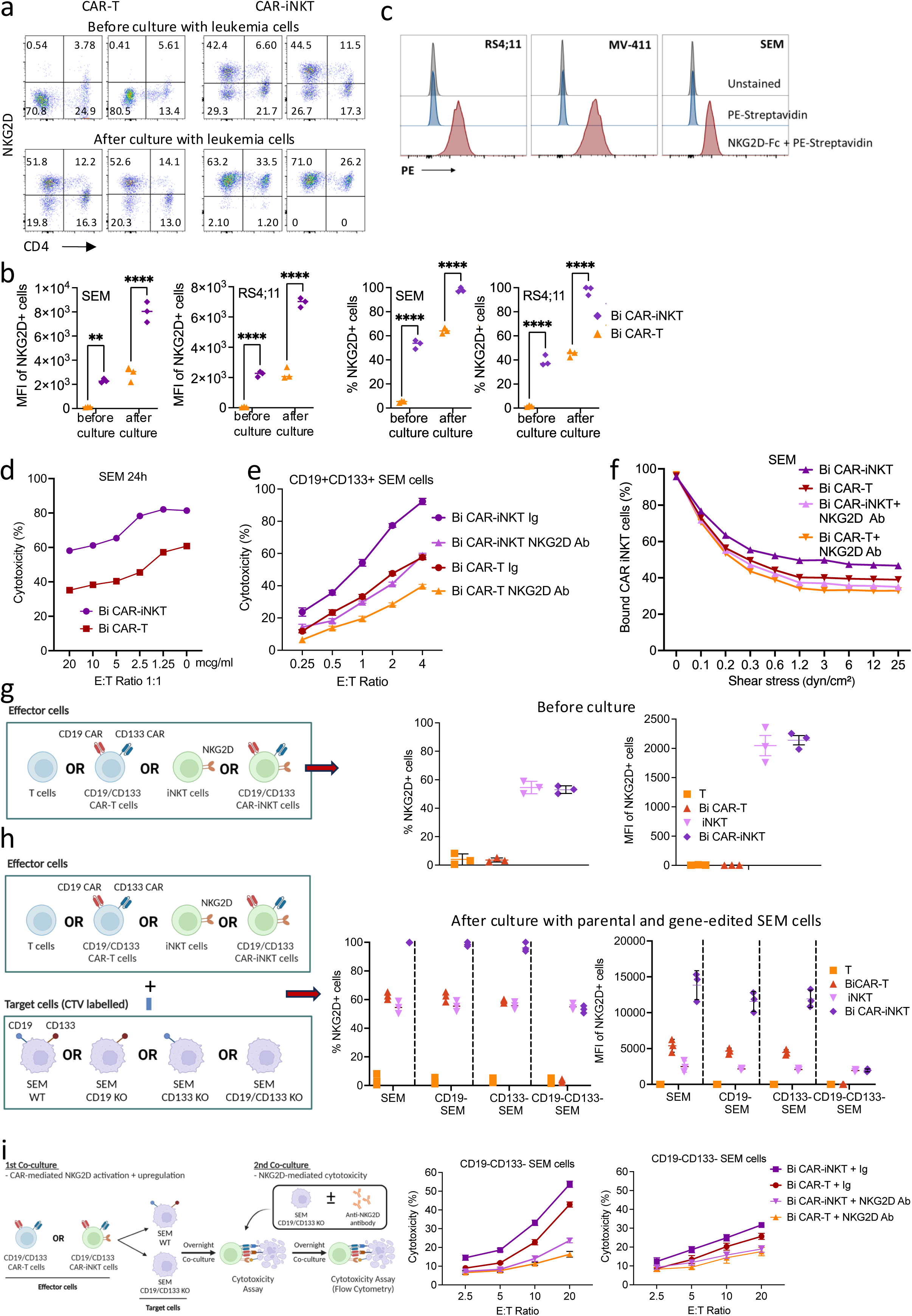
NKG2D-dependent anti-leukemic activity of CAR-iNKT. **a.** Flow-cytometric analysis of NKG2D expression on bi-specific CD19-CD133 CAR-T and -iNKT cells before and after overnight co-culture with SEM cells. **b.** NKG2D expression as measured by per cent of cells and mean fluorescence intensity in SEM and RS4;11 KMT2Ar cells. **c.** NKG2D ligand expression in the indicated KMT2A-r leukemia cell lines as assessed by staining with biotinylated NKG2D-Fc protein followed by fluorescent streptavidin**. d & e.** Cytotoxicity of bi-specific CAR-T and CAR-iNKT that had been pre-cultured with SEM cells against SEM cells in the presence of different concentrations (d) or 5μg of NKG2D mAb or Ig isotype control (e)**. f.** Avidity measurement of bi-specific CAR-iNKT vs CAR-T against SEM cells in the presence of NKG2D mAb or Ig isotype control. **g.** Left: Schematic of experiment. Right: NKG2D expression (% and MFI) in T/iNKT and CAR-T/-iNKT before co-culture with leukemia cells. **h.** Left: Schematic of experiment. Right: NKG2D expression after co-culture with the different SEM cell lines as shown. **i.** Left: Schematic of experiment. Middle: Cytotoxicity of bi-specific CAR-T and -iNKT that had been pre-cultured with CD19+CD133+ SEM cells against CD19-CD133-SEM cells in the presence of NKG2D mAb or Ig isotype control. Right: Cytotoxicity of bi-specific CAR-T and -iNKT that had been pre-cultured with CD19-CD133-SEM cells against CD19-CD133-SEM cells in the presence of NKG2D mAb or Ig isotype control. a-i: Data are from two independent experiments and two different iNKT donors.

To investigate the functional significance of these observations we tested the cytotoxic activity of bi-specific CAR-iNKT vs CAR-T that had been pre-exposed to leukemia cells, in the presence of an NKG2D blocking mAb or Ig control. We found a concentration-dependent decrease in the cytotoxic activity of CAR-iNKT and -T cells against the KMT2Ar SEM cell line in the presence of NKG2D mAb (**Figure 5d&e)** suggesting that the higher anti-leukemic activity of CAR-iNKT is largely NKG2D-mediated. Consistent with these results, the modestly higher avidity displayed by bi-specific CAR-iNKT when compared to CAR-T counterparts was also mitigated upon NKG2D blockade (**Figure 5f)**. Using NKG2D-Fc soluble protein to block the NKG2D ligands on target cells also abrogated the enhanced cytotoxicity of CAR-iNKT vs CAR-T thus further confirming the role of NKG2D in the differential reactivity of CAR-iNKT vs CAR-T against leukemia **(Suppl Figure 6b)**.

To investigate further the mechanism of upregulation of NKG2D expression, we co-cultured untransduced or CAR-transduced T and iNKT with SEM cells expressing CD19 only, CD133 only, both, or none (**Figure 5g&h)**. We found that expression of NKG2D in untransduced T and iNKT did not increase after exposure to CAR target-expressing or non-expressing SEM cells. By contrast, the higher baseline expression of NKG2D on CAR-iNKT than CAR-T commensurately increased in CAR-iNKT and CAR-T after their co-culture with CAR target-expressing cells i.e., CD19^+^CD133^+^, CD19^+^CD133^-^ or CD19^-^CD133^+^ SEM cells; however, upon co-culture with CD19^-^CD133^-^ SEM cells, both CAR-iNKT and CAR-T failed to upregulate NKG2D (**Figure 5h and Suppl Figure 6c)**. Together these results show that upregulation of NKG2D requires both expression of CARs and their engagement by their corresponding targets on leukemia cells.

Since bi-specific CAR-iNKT can eliminate CAR target high as well low-expressing leukemia cells *in vivo* (**Suppl Figure 2d-g),** we next tested whether CAR-iNKT could kill leukemia cells lacking complete expression of both CAR targets and the role of NKG2D in this process (**Figure 5i)**. We found that CAR-iNKT and to a lesser extent CAR-T cells that had been pre-exposed to CAR target-expressing leukemia cells, displayed higher cytotoxicity at high effector:target ratios against CAR target-negative SEM cells than CAR-iNKT and CAR-T that had been pre-exposed to leukemia cells lacking expression of the CAR targets (**Figure 5i)** and this effect was attenuated by NKG2D blockade. Similarly, CAR-iNKT pre-cultured with ^CRISPR^KMT2A-AFF1 ALL blasts upregulated NKG2D more efficiently than CAR-T (**Suppl Figure 6d)** and subsequently were able to kill CAR target-negative SEM cells more efficiently than CAR-T in an NKG2D-dependent manner (**Suppl Figure 6e).**

These findings highlight the ability of CAR-iNKT cells to enhance their innate, NKG2D-dependent effector functions in a CAR- and CAR target-dependent manner. Such a mechanism explains their higher efficacy compared to T cells and could help limit immune escape of CAR target-low/negative leukemia cells, potentially including those that might have undergone myeloid lineage switch, provided they express NKG2DL. In line with this, transcriptome analysis of paired diagnostic and relapse KMT2Ar ALL patient samples^39^ shows that while expression of *CD19* and *PROM1/CD133* is variably reduced upon lineage switch of B-ALL to myeloid blasts, leading to low-level expression of either CD19 or CD133, expression of NKG2D ligands and in particular of MICA-B remains the same or is higher in lineage switched myeloid blasts than in the paired presentation lymphoblasts **(Suppl Fig 6f)**.

### Ex vivo functional characterisation of CAR-iNKT cells at the single cell level

To gain insights into how the functional state of CAR-iNKT cells is altered *in vivo* upon their engagement with leukemia, we subjected pre-infusion CAR-iNKT and CAR-iNKT isolated at two timepoints from the bone marrow of CAR-iNKT-treated, leukemia-bearing or leukemia-free mice (i.e., five groups) to single cell TCRαβ and transcriptome analysis **(Figure 6a)**. Specifically, CAR-iNKT cells were isolated using CD2 immunomagnetic beads and pooled (from 3 mice) at two timepoints: on day 3 post CAR-iNKT treatment, when no leukemia cells were detected in the BM and spleen of treated animals and on day 15 when CAR-iNKT cells were still readily detectable **(Suppl Figure 7a).**

**Figure 6.**
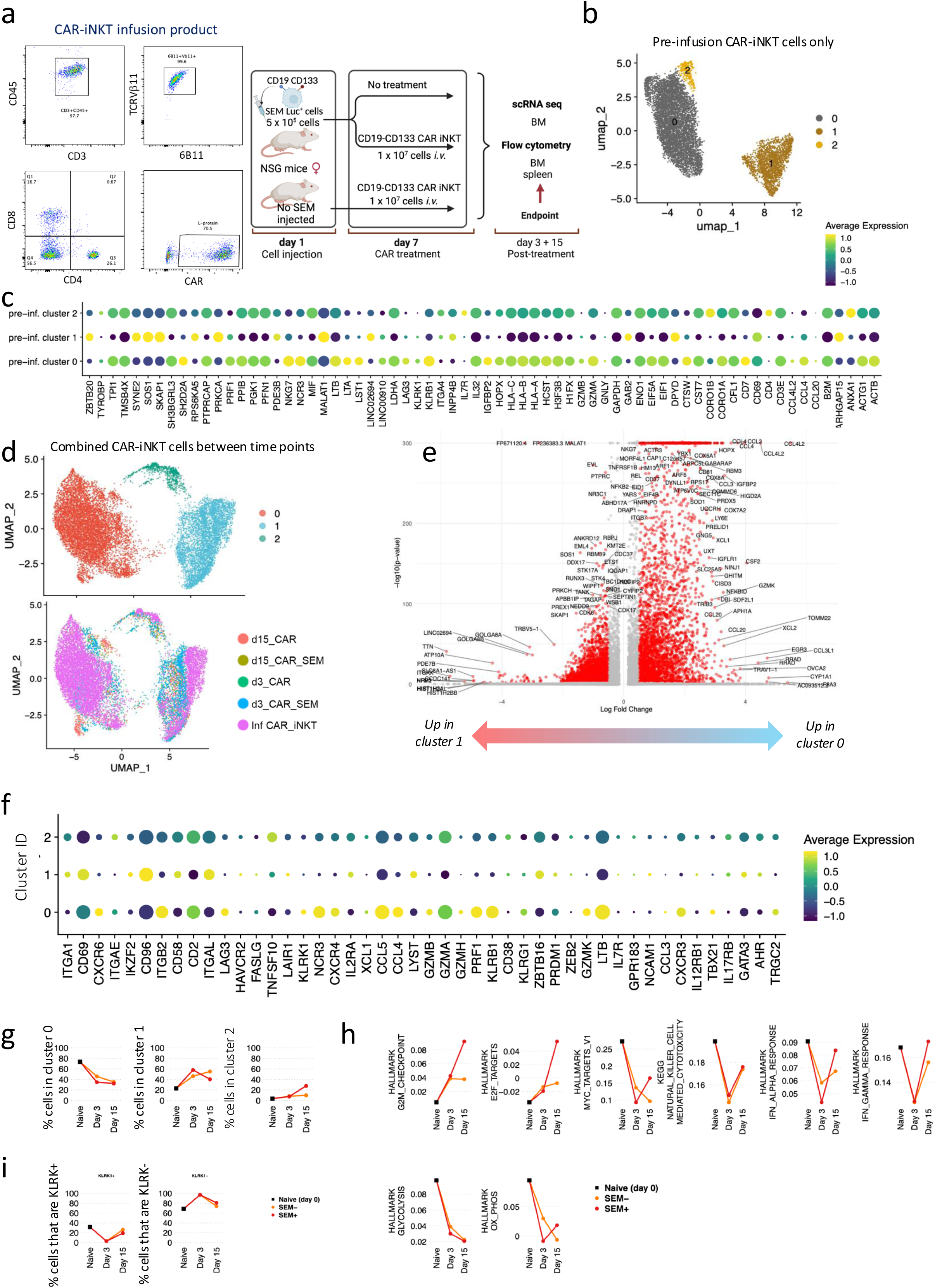
Ex vivo transcriptome analysis of bi-specific CAR-iNKT cells. **a.** Immunophenotypic profiling of pre-infusion CAR-iNKT cells and design of the experiment. **b.** UMAP embedding of gene expression data of the pre-infusion CAR-iNKT. **c.** Bubble plot showing comparative expression levels of genes of immunological relevance in clusters 0,1&2 from b. **d.** UMAP embedding of gene expression data of all experimental groups after batch correction coloured by cluster ID (top) and by sample (bottom). **e.** Volcano plot showing differential gene expression between clusters 0 & 1 (log2FC>1 and padj< 0.05). **f.** Bubble plot showing comparative expression levels of genes of immunological relevance in clusters 0, 1 and 2 from d. **g.** Frequency of cells in different clusters over the time course of the experiment. **h**. Dynamic variation in the score of indicated enriched pathways between the pre-infusion product (naïve) and day 3&15 CAR-iNKT. All pathways are significantly different between samples (p-value <1e-10). **i.** Dynamic variation in the proportion of KLRK1(NKGD)+/- cells between the pre-infusion product and day 3&15 CAR-iNKT.

First, single cell gene expression-based dimensionality reduction of the pre-infused CAR-iNKT cells identified three clusters (C0-2) with, as expected, the minority of cells expressing CD4 and co-localising mostly in cluster 2 **(Figure 6b)**. Differential gene expression (DGE) between CD4+ vs CD4-CAR-iNKT **(Suppl Table 1)** identified the latter as enriched in a cytotoxic transcriptional signature (*PRF1*, *NKG7*, *NCR3*, *LTA*, *LTB*, *KLRK1*(NKG2D), *KLRB1* (CD161), *GZMB/A*), consistent with their known biology **(Figure 6C)**.

Single cell gene expression and TCR expression-based dimensionality reduction of all CAR-iNKT cells identified three clusters of cells (C0, C1 & C2) with all five groups represented in each cluster **(Figure 6d).** As expected, although not captured in all cells, the invariant TCRAV10-TRAJ18 sequence was by far the most dominant TRA clonotype **(Suppl Figure 7b-d)**.

Focusing on genes over-expressed in cluster 0 vs 1 (n=1642, Log_2_FC > 1 & Padj< 0.05**; (Figure 6e** and **Suppl Table 2)** revealed enrichment for oxidative phosphorylation and enhanced protein synthetic capacity (Myc Targets V1; MTORC1 signalling; **Suppl Figure 7e)** suggesting that in a sizeable fraction of CAR-iNKT favourable energy production and protein anabolic pathways support their function. At a single gene expression level, cluster 0, compared to clusters 1&2 emerged with a robust profile of cytotoxic potential with perforin (*PRF1*), several granzyme genes (GZMB, GZMH, GZMK) and activating NK cell receptor genes (*KLRK1/NKG2D, NCR3, LTB*) preferentially and highly expressed in cluster 0 cells (**Figure 6f)**. Of note, with the exception of *GZMH* and *GZMK* the expression of which was reduced, cells retained their expression of *GZMA*, *GZMB* and *PRF1* after 3 and 15 days **(Suppl Figure 7f**).

DGE analysis between the day 3 groups revealed only a handful of variable genes, whereas a larger number of variable genes were identified between day 15 groups **(Suppl Figure 7g and Suppl Table 2**). Day 15 iNKT cells from leukemia-bearing mice over-expressed 264 genes, compared to iNKT from leukemia-free mice (log2FC>1, padj<0.05), with the most enriched pathways amongst over-expressed genes corresponding to cell cycle (G2M checkpoint, E2F targets) dominating **(Suppl Figure 7e**), thus highlighting the fact that >10 days after clearing leukemia CAR-iNKT retain robust proliferative potential. Applying a transcriptional score for the cell cycle S phase in CAR-iNKT from day 15 leukemia-bearing vs leukemia-free mice showed the former to have a higher proliferative status (P<6.2×10^-6^) while no differences were found for day 3 counterparts **(Suppl Figure 7h).** In line with this, while frequency of C0 declined, frequency of cells in cluster 2, which is the most proliferative of the three clusters **(Suppl Figure 7i)** increased on day 15 only in leukemia-bearing but not leukemia free mice **(Figure 6g).**

To obtain a dynamic overview of the function of CAR-iNKT we scored pathways enriched between the five experimental groups (P-value <1^-10^) and plotted them against the pre-infusion, day 3 and 15 timepoints **(Figure 6h).** We found that proliferation increases *in vivo* with time, with a higher score in CAR-iNKT from leukemia-bearing mice (G2M checkpoint, E2F targets). While on day 3 cytotoxic and IFN (IFNα & γ) responses are blunted, in day 15 iNKT cells they are both restored, with IFNα & γ responses scoring higher in CAR-iNKT from leukemia-bearing mice. This is mirrored by a reduction in metabolic pathways including glycolysis, but relative preservation of oxidative phosphorylation in day 15 iNKT cells from leukemia-bearing mice.

Finaly, scoring the frequency of KLRK1/NKG2D-expressing cells shows that after their decline on day 3 they recover on day 15 **(Figure 6i).**

Together these findings suggest that CAR-iNKT retain and enhance their effector and proliferative functions after their *in vivo* targeting and eradication of leukemia. To corroborate this prediction we isolated iNKT cells from the BM of bi-specific CAR-iNKT-treated, leukemia-bearing animals shown in Fig 2a&b involving a different CAR-iNKT donor. We found presence of iNKT cells in the BM of all treated animals treated with 10^6^ cells on day 6 **(Suppl Fig 8a)**, isolated them and investigated their functional properties *in vitro*. Explanted CAR-iNKT cells proliferated robustly in the presence of αGalCer (>100-fold over two weeks; **Suppl Fig 8b**), retained the same level of CAR expression as the originally injected cells (**Suppl Fig 8c**), and following expansion they retained their ability to exert a powerful cytotoxic effect when cultured with CD19^hi^CD133^hi^ or CD19^lo^CD133^lo^ SEM leukemia cells **(Suppl Fig 8d)**. When tested against the CD19-expressing C1R and C1R-CD1d B cells with or without αGalCer, explanted bi-specific CAR-iNKT cells exerted the highest cytotoxicity against αGalCer-pulsed C1R-CD1d cells **(Suppl Fig 8e)**, thus confirming that their iTCR-CD1d axis is functionally preserved and can contribute to the anti-tumor function of CAR-iNKT cells. Therefore, as predicted by transcriptome analysis, even after 4 weeks of pre infusion *in vitro* expansion, ∼5 weeks of *in vivo* persistence in leukemia-bearing mice, and another 3 weeks of ex vivo expansion, bi-specific CAR-iNKT cell retain robust proliferative and cytolytic anti-leukemia activity.

### Impact of bi-specific CAR-iNKT cells on normal human hematopoiesis

Given the expression of CD133 by a proportion of normal primitive HSPC **(Suppl Fig 1d)**, we used a humanised xenograft model to assess potential off-tumor effects of CD19-CD133 bi-specific CAR-iNKT by transplanting cord blood (CB) CD34+ cells into sub-lethally irradiated NSG-SGM3 mice **(Suppl Fig 9a)**. In two independent experiments, engrafted NSG-SGM3 mice (confirmed by hCD45 expression in >1% of PB cells; **Suppl Fig 9b and Suppl Fig 10a)** were treated with either PBS or 10^7^ CD19-CD133 bi-specific CAR iNKT cells. On-target, off-leukemia activity of CAR iNKT cells was confirmed by a significant reduction in the frequency of CD19+ B cells in the PB at day 1 and day 3 post CAR-iNKT injection, compared to controls (**Suppl Fig 9c, Suppl Fig 10b**). Long-term engraftment in PB (18 weeks) or BM (at cull 18-21 weeks post-transplantation) showed no significant difference between CAR iNKT- and PBS-treated mice (**Suppl Fig 9b&d, Suppl Fig 10a,c)**, either in total hCD45+ cells or, in BM, in the frequencies of B, T, myeloid or CD34+ cells within the hCD45 compartment (median values for B cells: 25.7% vs 26.1%; T cells: 22.4% vs. 25.8%, myeloid cells: 43.5% vs. 46.4% and CD34: 3.2% vs. 2.3% of hCD45 respectively; **Suppl Fig 9d)**. This confirms comparable multi-lineage long term human hematopoietic reconstitution in PBS- and CAR iNKT-treated mice, suggesting there is no significant CAR iNKT-mediated toxicity against primitive HSPC.

## Discussion

For immunotherapy of cancer to be effective, therapeutic strategies need to be specifically tailored to the biology of each individual disease. A combination of chemoresistance and a high risk of CNS disease contribute to the adverse prognosis of KMT2Ar ALL, including infant ALL. Here we deploy allogeneic iNKT cells, ‘innately’ more powerful effectors than T cells, and equip them with bi-specific CD19-CD133-targeting CARs that engage co-expressed leukemia-associated antigens in this disease. We show that compared to mono-specific counterparts and bi-specific CAR-T, CD19-CD133 bi-specific CAR-iNKT not only have a more potent anti-leukemia activity in faithful and clinically relevant models of KMT2Ar-ALL, they are also able to eradicate medullary and leptomeningeal leukemia with variable CAR target expression and to induce sustained remissions without discernible hematologic toxicity.

In all cases, enhanced avidity of bi-specific CAR-iNKT, in line with recent work^19,40,41^, correlated with *in vivo* efficacy. Increased avidity of bi-specific over mono-specific CAR-iNKT would be particularly important for the effective targeting of CAR antigen-low disease, as we demonstrate in one of our *in vivo* models, and would potentially mitigate subsequent CAR target-low relapse often seen in KMT2Ar-ALL and other forms of ALL treated with CD19 CAR-T^8,11^.

We also show that higher and dynamic expression of NKG2D by CAR-iNKT plays an important role in their higher avidity. Previous work demonstrated the importance of NKG2D expression by CD4^-^ iNKT and by CAR-iNKT for enhancing iNKT activation independent of the iTCR-CD1d interaction when engaged by NKG2D stress ligands^35,42^. Here, we demonstrate a novel process of dynamic NKG2D expression by both CD4^-^ and CD4^+^ iNKT subsets. Importantly, this allows targeting of leukemia cells with variable expression of CAR targets, including cells lacking expression of both CAR targets, in an NKG2D-dependent manner. These findings are analogous to TCR-MHC class I-dependent upregulation of NKG2D in conventional CD8^+^ T cells, allowing them to target tumor cells with loss of MHC molecules in a NKG2D-dependent manner^43^. This mechanism is more powerful in CAR-iNKT than CAR-T and is important not only for eradication of leukemia with low expression of one of the CAR targets, but may also reduce the risk of lineage switch by directly targeting lineage-switched blasts which characteristically exhibit partial or complete loss of CD19 and CD133 expression^39^. As expression of NKG2D ligands in lineage-switched blasts remains high or even higher than in the original ALL cells, NKG2D-mediated anti-leukemia activity of CAR-iNKT cells is potentially attractive as a strategy for targeting lineage-switched cells even at an early stage. Further studies will be needed to directly investigate these predictions *in vivo*.

Our study clearly showed the curative potential of CAR-iNKT against different levels of leukemia burden, from low to high, but only when used at higher dose levels *in vivo*. Similarly, the *in vitro* NKG2D-mediated anti-leukemia activity against CAR target-negative leukemia was also more evident at higher E:T ratios. Importantly however, although the superior anti-leukemic activity of CAR-iNKT over CAR-T was evident at limiting dose levels, this did not translate into curative potential despite their longer persistence and maintained functionality. This is in line with recent pre-clinical data showing that failure of immunotherapy is often not due to tumor-intrinsic mechanisms of immune escape but simply due to insufficient numbers of effector cells reaching all tumor sites when relatively low therapeutic doses are used^44^. These findings should inform clinical development and call for use of high doses of CAR-iNKT in future clinical trials.

A major advantage of higher bi-specific CAR-iNKT cell doses is their astonishing ability to confer protection from, and eradication of established meningeal leukemia, which affects a higher proportion of infants with KMT2Ar ALL^45,46^. Delineating the mechanism(s) of this critical property of iNKT cells, which we previously reported in brain lymphoma^26^, is likely to have broader therapeutic implications for leptomeningeal metastases in a variety of cancers, including breast cancer, melanoma and lung cancers^47^, while in the case of ALL, CAR-iNKT may be an alternative to toxic intrathecal methotrexate for CNS prophylaxis. Our previous transcriptome analysis suggested that expression of VLA-4^26^, an integrin required for endothelial adhesion and crossing of the blood brain and choroid plexus-brain barriers, was higher in CAR-iNKT than in CAR-T^26^.

Our transcriptional analysis of CAR-iNKT cells shows that gene expression programmes associated with cytotoxic and proliferative potential as well as metabolic fitness that are present in the pre-infusion CAR-iNKT cells, are preserved at two weeks following their *in vivo* interaction with leukemia cells. These findings are in line with the fact that explanted CAR-iNKT retain their ability to proliferate and exert robust cytotoxicity against leukemia cells.

We propose that future clinical application would likely employ allogeneically-sourced, bi-specific CAR-iNKT as a swift bridging therapy to induce remissions in relapsed/refractory KMT2Ar-ALL prior to allo-HSCT^48^. This approach may have the added benefit of eradicating CD19-KMT2Ar leukemic stem/progenitors that may drive relapse and/or lineage switch^9,39,49^. While not as developed as other forms of cellular immunotherapy such as CAR-T and CAR-NK cells, allogeneic CAR-iNKT and iNKT-based immunotherapeutics are at the verge of clinical development with early clinical results suggesting high efficacy with, as predicted, lack of aGVHD and importantly, mild or no cytokine release syndrome and neurotoxicity^42,50,51^. Importantly, our study showed that at dose levels that induce B cell depletion, bi-specific CAR-iNKT did not significantly impact long term hematopoiesis, in line with a clinical trial of autologous CD133 CAR-T in patients with solid tumors which reported only transient grade 2 hematologic toxicity^52^.

We conclude that bi-specific CAR-iNKT cell immunotherapy is a very effective treatment for pre-clinical aggressive KMT2Ar-ALL, it outperforms CAR-T cell immunotherapy in an NKG2D-dependent manner, it eradicated leptomeningeal leukemia and has the potential to protect from immune escape with CAR target antigen-low disease and potentially lineage switch without discernible hematologic toxicity. These findings provide the basis for clinical development of bi-specific CD19-CD133 CAR-iNKT cells as an ‘off-the-shelf’ treatment for KMT2Ar-ALL. More broadly, the ‘innate’ ability of CAR-iNKT cells to effectively target CAR target-negative/low as well as CAR target-expressing leukemia cells calls for their development for a wide range of high-risk malignancies with leptomeningeal involvement.

## Methods

### Primary human samples

#### iNKT and T cells

Peripheral blood mononuclear cells (PBMCs) obtained from healthy donors were isolated by density gradient centrifugation and used as a source of CD3+ lymphoid cells for CAR engineering.

#### Human fetal hematopoietic stem and progenitor cells

These were provided for the purposes of this research by the Human Developmental Biology Resource (HDBR, www.hdbr.org), regulated by the UK Human Tissue Authority (HTA, www.hta.gov.uk) and covered under ethics granted by NHS Health Research Authorities: North East – Newcastle & North Tyneside Research Ethics Committee (REC: 23/NE/0135) and London - Fulham Research Ethics Committee (23/LO/0312). Informed consent was obtained from all participants, who donated human fetal tissue for research without receiving any monetary compensation. FL samples used for CRISPR/Cas9 KMT2A-AFF1 translocation experiments underwent CD34 magnetic bead selection at the time of sample processing and were cryopreserved for future use as described^53^. Cord blood samples were obtained from NHSBT under ethical approval (REC: 21/LO/0195) and CD34 cells were selected as above.

#### Leukemia samples

ALL patient samples were obtained from VIVO Biobank, UK after appropriate review of our research project to ensure that it was covered under their ethics approval granted by NHS HRA South West - Central Bristol Research Ethics Committee (REC: 23/EM/0130). Two infant KMT2Ar ALL samples were obtained from Our Lady’s Children’s Hospital, Crumlin, Dublin, Ireland and one from Oxford University Hospital Trust under ethical approval (REC: 21/LO/0195). Informed consent was obtained from all participants or those with parental responsibility, and participants did not receive any monetary compensation. Infant and paediatric KMT2Ar ALL samples from patients being treated at Great Ormond Street Hospital for Children, London were analysed by flow cytometry as part of their diagnostic workup after informed consent. All patient samples/data were anonymized at source, assigned a unique study number and linked. Patient derived xenografts: KMT2Ar ALL PDX cells were provided by the Halsey lab (Glasgow).

Model cells of infant B-acute lymphoblastic leukemia were derived by introducing a *KMT2A -AFF1* translocation into CD34 selected human fetal liver cells using CRISPR and transplanted into NSG mice for *in vivo* expansion (^CRISPR^KMT2A-AFF1 ALL) as described in ^32^.

### Cell lines

SEM, RS4;11 and KOPN8 cell lines were available in the Milne lab. SEM cell line was further modified in the following ways: SEM cells were transduced with non-replicative MIGR1 retrovirus to co-express Luciferase and dsRed as an expression marker. All cell lines were purchased and maintained under recommended conditions. SEM cells were cultured in IMDM (Gibco) supplemented with 10% FCS. RS4;11 and KOPN-8 cells were cultured in RPMI (Gibco) supplemented with 10% FCS.

### CRISPR/Cas9 gene editing of SEM cells

*CD19* and *PROM1* knockouts were performed in SEM cells using pools of targeted CRISPR RNP guides purchased from Synthego (**Suppl Table 3**) and following the recommended procedures and concentrations for the Neon Transfection System (Invitrogen: MPK1096) and Cas9 Nuclease V3 (IDT: 1081059)^32^. Knockout cells recovered in standard culture conditions for 3-7 days post-transfection before immunophenotyping was performed to select for complete knockout before continuing on to additional experiments.

### CAR constructs

For the anti-CD19 single-target CAR, we employed the FMC63 antibody clone, fused with the hinge, transmembrane, and partial intracellular domain of CD8α (amino acids 128-210 of CD8α), followed by the CD28 co-stimulatory domain and CD3ζ activation domain. The anti-CD133 single-target CAR was constructed using the AC133 antibody clone, incorporating the same CD8α components as the anti-CD19 CAR, but coupled with the 4-1BB co-stimulatory domain and CD3ζ activation domain. The bi-specific CD19-CD133 dual CAR was engineered by linking the aforementioned single-target CARs via a T2A peptide sequence. The schematic structures of these CAR constructs are illustrated in Figure 1a. Following synthesis of the complete coding sequences, these constructs were cloned into the pSIEW lentiviral transfer plasmid using appropriate restriction enzyme digestion and ligation approaches.

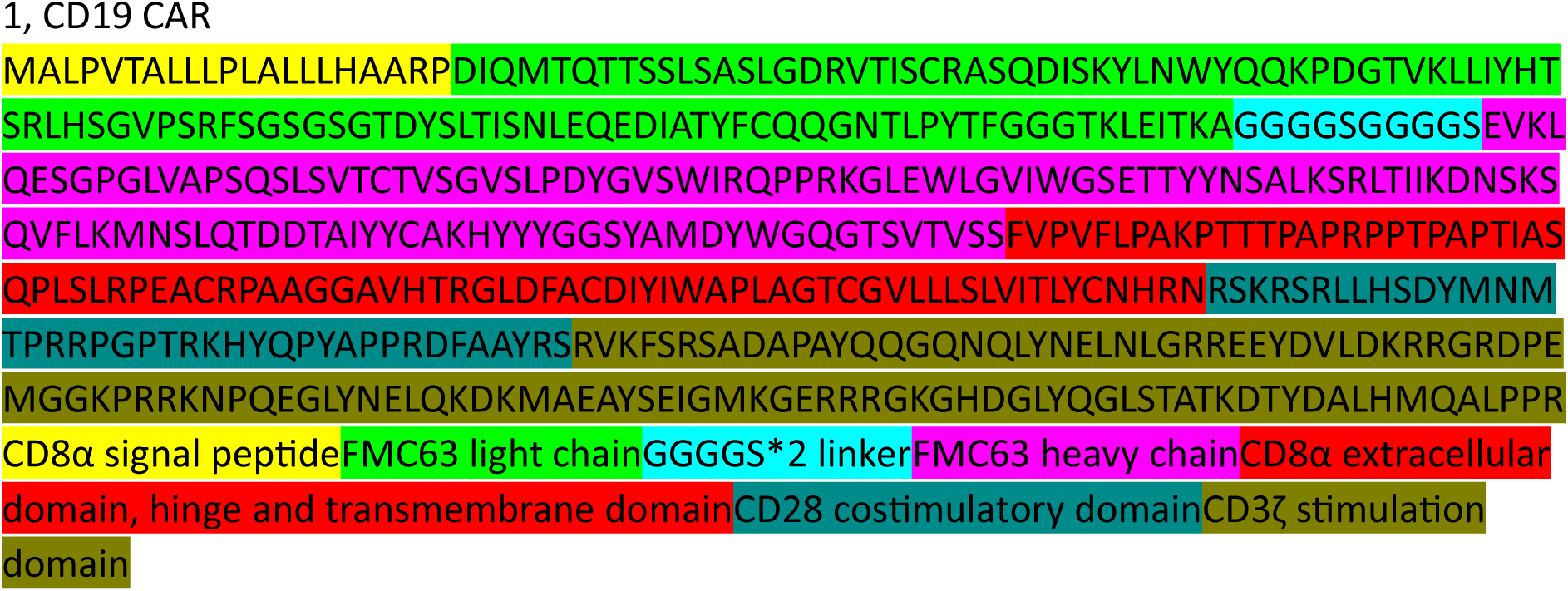

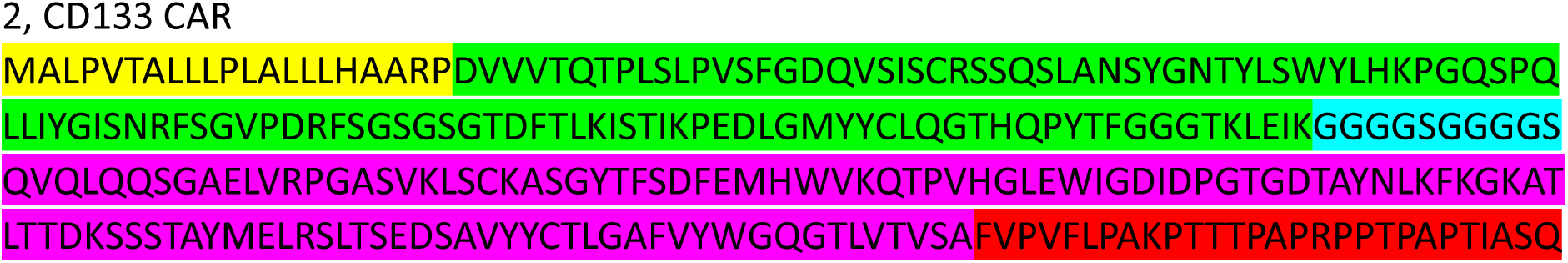
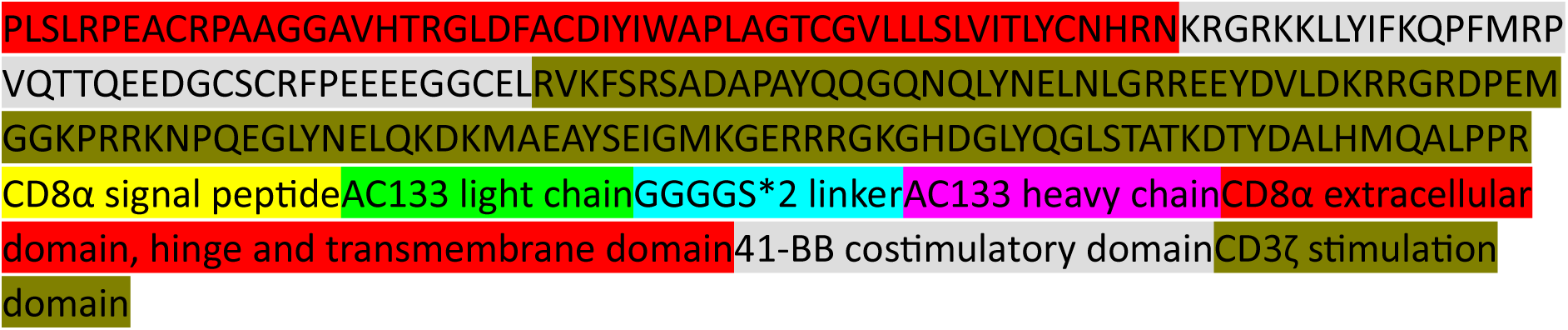

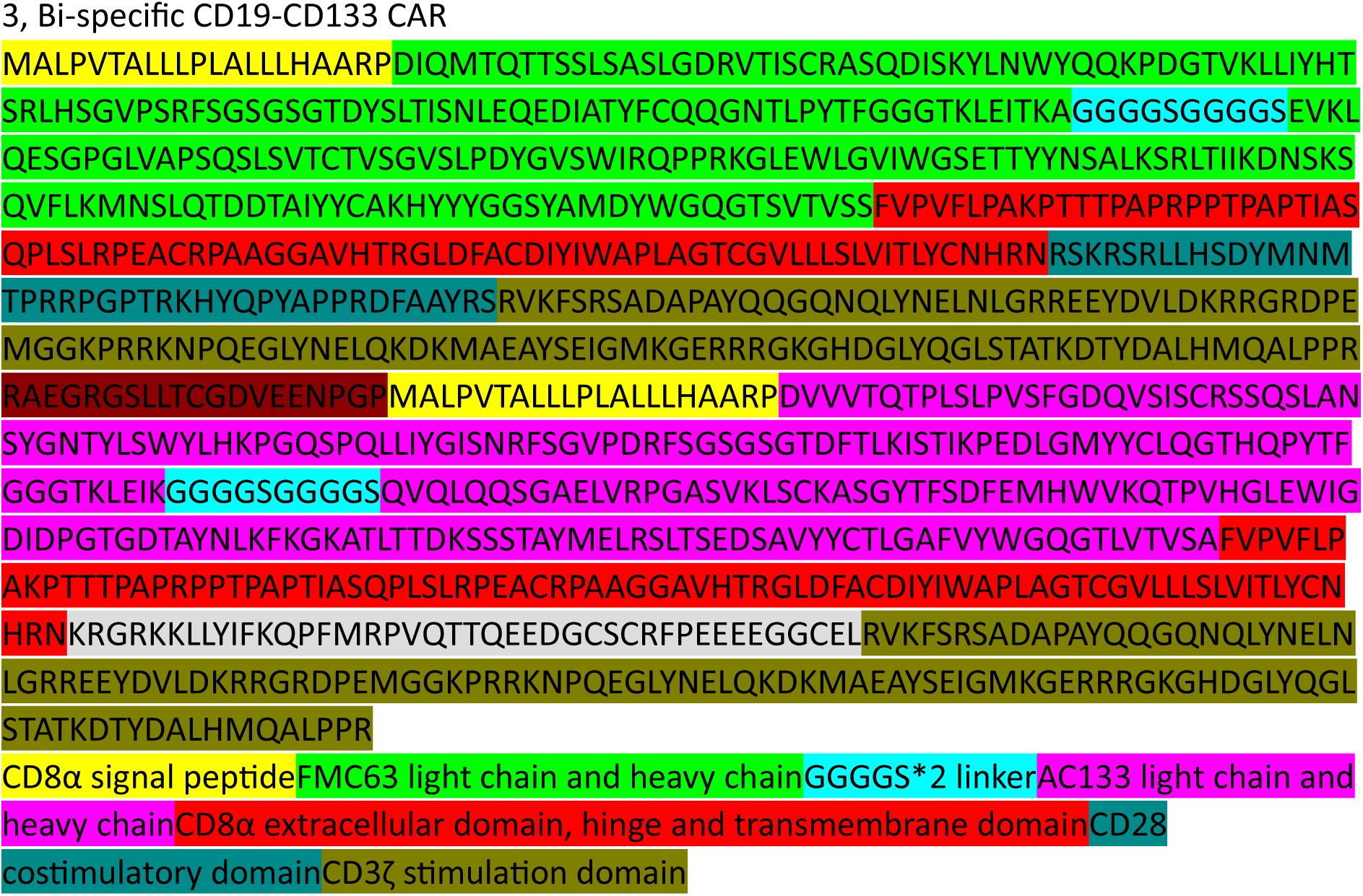

### CAR-iNKT and CAR-T generation

TCRVα24Jα18+ lymphocytes were immunomagnetically sorted from PBMC using anti-human iNKT cell microbeads (Miltenyi Biotec). Purified iNKT cells were seeded in 24- or 48-well plates at a 1:1 ratio with irradiated (3500 rad) autologous mononuclear cells (iAPC) and activated with Dynabeads Human T-Activator CD3/CD28 (Gibco™) at a 1:1 beads-to-cell ratio in T cell medium at a density of 1-5 × 10^4^ cells per ml. IL-15 (Miltenyi Biotec) at 30 IU/ml and 150 IU/ml was added at the time of seeding and 12 hours later, respectively. Within 48 hours, activated iNKT cells were transduced with concentrated CAR lentivirus using an MOI of 2-5 in the presence of 8 μg/ml pre-coated retronectin, with spinoculation for 90 minutes at 1000G. After 8-12 hours, cells were resuspended in fresh medium supplemented with 150 IU/ml of IL-15 and allowed to rest for 4 days before assessment of viability and CAR expression. CAR transduction efficiency was determined by flow cytometry as the percentage of L-protein+ cells relative to untransduced controls. CAR+ cells were re-stimulated with a 1:1 ratio of irradiated C1R-CD1d cells loaded with αGalCer (100 ng/ml), IL-15 (30 IU/ml), and with an additional 150 IU/ml of IL-15 added 12-24 hours later.

Subsequently, cells were expanded for 14 to 35 days, assessed for purity, and used for in vitro assays. Alternatively, CAR iNKT cells were harvested during the exponential growth phase, cryopreserved in 10% DMSO, and stored in liquid nitrogen until use. Untransduced iNKT were generated in the same way, with the omission of lentiviral transduction step. CAR-T were generated as previously described^26,27^.

### *In vitro* assays

#### Flow cytometry

Cells were stained with fluorophore-conjugated monoclonal antibodies in PBS with 2% FBS and 1mM EDTA for 30 minutes and analyzed via LSR Fortessa X50 or FACS sorted via BD Aria instruments using BD FACSDiva software (v8.0.2). Antibodies used are detailed in **Suppl Table 4**. Flow cytometry antibodies were validated by titration in-house using primary human fetal mononuclear cells (MNC) or NSG mouse BM. Analysis was performed using FlowJo software (v10.7.1) where gates were set using unstained and fluorescence minus one (FMO) controls.

#### Cytotoxicity assays

These were performed as previously described^26^. Briefly, CellTrace™ Violet (Invitrogen)-labelled targets were incubated at the indicated ratios with effector cells for 4 or 16-24 hours. As controls, targets and effectors alone were simultaneously incubated to determine spontaneous cell death. Cells were then harvested and 7-AAD was added prior to flow cytometric analysis on BD LSR Fortessa Flow Cytometer, using BD FACSDiva software version 6.0. Specific cytotoxic activity was determined as ((% sample (7-AAD+, Violet+) − % spontaneous (7-AAD+, Violet+)) / (100 - %spontaneous (7-AAD+, Violet+)) × 100. All assays were run in duplicates or triplicates and analysed using FlowJo 10.9.0.

Intracellular cytokine expression assays were performed as previously described^26,27^.

### Flow-chamber avidity

A µ-VI 0.4-A 6 channel slide (Ibidi) and a pump power system were used for the cell-binding avidity assay. Cell lines or primary leukemia cells, were attached to poly-L-lysine-coated chips as a monolayer for at least 3 hours prior to testing. CellTrace™ Far Red -labelled (Invitrogen) target cells were allowed to bind for 5 minutes before the flow rate and pressure were ramped up. The flow rate was increased from 0 to 25.6 ml/min of flow through the chamber, with real-time imaging using a EVOS M5000 Imaging System (Thermo Fisher) conducted under both static and flow conditions. Counting of target cells remaining attached at each flow rate was analysed using ImageJ software.

### *In vivo* experiments

#### Animals

All experiments were performed under two separate project licenses approved by the UK Home Office under the Animal (Scientific Procedures) Act 1986 after approval by the Oxford and Imperial College Animal Welfare and Ethical Review Bodies; and in accordance with the principles of 3Rs (replacement, reduction and refinement) in animal research.

Experimental animals were 6–8-week-old female NOD.Cg-PrkdcscidIl2rgtm1Wjl/SzJ (NSG) mice or 6-week-old female NSGS mice. Male mice were used for some secondary and tertiary xenotransplantation experiments. Mice were housed in IVC cages, and kept at a 12-hour light/dark cycle, 21-22°C temperature and 45-65% relative humidity. They had red tunnels or houses and balconies in the cages as enrichment.

### Bioluminescence Imaging (BLI)

BLI were collected on an IVIS Lumina XR III Imaging System using Living Image software (PerkinElmer). Mice were anesthetized and maintained under inhalational anaesthesia via a nose cone with 2% isoflurane (Zoetis UK)/medical oxygen. A single intraperitoneal (IP) injection of 150 mg/kg D-luciferin (Goldbio) in PBS was administered to all mice 10 minutes before scanning. Up to three mice were imaged simultaneously in a 12.5 cm field of view (FOV) with a minimum target count of 30,000 and exposure times ranging from 0.5 to 3 minutes at medium binning, with additional images acquired at low binning levels to maximize sensitivity and spatial resolution where required. Both ventral and dorsal scans were acquired for each mouse. The dorsal and ventral signals were quantitated separately through region of interest (ROI) analysis using Living Image software (Aura-4.0.8) and expressed in radiance (units of photons/sec) as a total signal summation normalized to the ROI area. Where required, normalized background signal from similarly sized ROIs was subtracted

### Leukemia Models

#### SEM model

Six-week-old NOD/SCID/IL-2Rγ-null (NSG) female mice (Charles River, UK) were handled in accordance with the 1986 Animal Scientific Procedures Act and under a United Kingdom Government Home Office–approved project licence PP8553679. The animals were housed at the Hammersmith Central Biomedical Services (CBS) facility, Imperial College London. On day 1, all animals were injected with 5 × 10^6 or 1 × 10^6 luciferase-expressing SEM cells via the tail vein (*iv*), followed by bioluminescence imaging (BLI) monitoring on day 6 to confirm engraftment. On day 7, day 12, or day 16, the mice were randomized to either no treatment or immunotherapy with CAR-T or CAR-iNKT cells generated from the same donor. Thereafter, BLI was performed twice a week until day 21 and weekly until the end of the experiment. All mice were euthanized according to protocol when either experimental or humane endpoints were reached.

#### ^CRISPR^KMT2A-AF4 ALL model

As with the SEM cells, on day 1, all animals were injected with 1 × 10^6 *^CRISPR^KMT2A-AF4* cells via the tail vein (iv), followed by tail vein blood collection to determine HLA-ABC or CD19, CD133 expression by flow cytometry on day 6 to confirm engraftment. On day 7, the mice were randomized to either no treatment or immunotherapy with CAR-T or CAR-iNKT cell generated from the same donor. All mice were euthanized according to protocol when either experimental or humane endpoints were reached.

### CAR-iNKT hematological toxicity assays in humanised mice

All experiments were carried out under a United Kingdom Government Home Office–approved project licence PP2666723. 7–9-week-old NSGS mice (n=12) were sub-lethally irradiated with two doses of 1.25Gy six hours apart (2.5Gy total) and injected via the tail vein with 60,000 cord blood CD34+ cells. Engraftment was monitored by peripheral blood sampling every 3 weeks. Engrafted mice (>1% human CD45 cells in peripheral blood), were divided into control group (received PBS only) or treatment group, treated with 10 million bi-specific CD19-CD133 CAR-iNKT cells at either 9 weeks or 15 weeks post CB CD34+ transplantation. Additional blood samples were taken at D+1 and D+3 post CAR-iNKT injection. Animals were monitored regularly using a standardized physical scoring system, and any mouse found to be in distress was humanely killed. All well mice were culled between 15-21 weeks post CB transplantation to assess long term bone marrow engraftment. Bone marrow was harvested from these mice for analysis.

### Brain histopathology

Murine heads were stripped of soft tissues, fixed in 10% neutral-buffered formalin (CellPath) and decalcified in Hilleman and Lee EDTA solution (5.5% EDTA in 10% formalin) for 2–3 weeks. Following paraffin embedding, hematoxylin and eosin staining (Sigma-Aldrich) was performed on 5-mm brain sections. Anti-CD19 immunohistochemistry on paraffin-embedded sections was performed as previously described^54^.

Imaging used Axiostar Plus or Axio Imager M2 microscopes with AxioVision and ZEN software (Carl Zeiss, Cambridge, United Kingdom). CNS infiltration was assessed by an experienced pathologist (Halsey) blinded to treatment allocation. 5-6 coronal slices and 2 blocks per head were examined and each block was assigned a score from 0= no infiltrate seen, 1= scattered occasional cells, 2= mild infiltrate, 3=moderate infiltrate, 4= heavy infiltrate.

### Single cell transcriptome-TCR combined analysis

#### Experimental design

Bone marrows from either day 3 or day 15 post-CAR-iNKT injection were pooled from 3 NSG mice per group of mice that either did or did not receive SEM cells, before CAR-iNKT enrichment and purification.

#### Ex vivo selection of iNKT

CAR-iNKT cells were enriched by magnetic bead selection following the manufacturers protocol against human CD2 (Miltenyi 130-091-114). CD2 positive cells were then further purified by FACS sorting staining using Labelling Check Reagent (Miltenyi 130-122-219) and 7AAD viability stain (Cayman 11397) with additional antibodies against mouse-CD45, human-CD45, and human CD19. The number of CD2+ cells recovered were 2768 to 7645 per pool (from 3 mice). In addition, 30,000 pre infusion CAR-iNKT cells from the same donor were also FACS sorted for single cell analysis. We then immediately proceeded with the TCR/RNAseq protocol for the sorted cells (10x genomics CG000331 Rev C).

### Single Cell Multi-omics sequencing and pre-processing

scRNAseq transcriptome processing was performed using the Chromium 10x system involving GEM generation, post GEM-generation clean-up, cDNA amplification and DNA quantification. The library was sequenced using the Illumina NovaSeq platform. Chromium Single Cell Immune Profiling Reagent Kits v1.1 solution was used to deliver a scalable microfluidic platform for gene expression (GEX) and VDJ TCR profiling. Libraries were generated and sequenced from the cDNAs and 10x Barcodes were used to associate individual reads back to the individual partitions.

The analysis pipeline applied to process Chromium single-cell data to align reads and generate feature-barcode matrices was performed as previously described^55^. The raw FASTQ files from scRNAseq and TCR sequencing were processed using Cellranger v7.1.0 in two separate steps. Briefly, gene expression FASTQ files were processed using Cellranger count to perform alignment, filtering, barcode counting, and UMI counting, using 10X Genomics’ GRCh38 v2020-A reference. For the TCR V(D)J analysis ‘cellranger vdj’ command was used taking GRCh38-alts-ensembl-7.1.0 as the reference to align, assemble and annotate T cell receptor sequence.

### Filtering, doublet detection and batch correction

For each sample, cells with fewer than 500 transcripts or 500 genes of >15% mitochondrial genes were filtered out. Normalisation and scaling were done using the standard Seurat pipeline. Principal component analysis (PCA) was performed on 10,000 highly variable genes (HVGs), excluding highly variable genes encoding for TCR variable chain, ribosomal proteins, heat shock proteins, mitochondrial proteins, cell cycle proteins, HLA, and noise-related genes (MALAT1, JCHAIN, XIST). These were used to compute 50 principal components, then *Harmony* was performed for batch correction^56^, UMAP for dimensionality reduction, and the Louvain algorithm was used for clustering (resolution: 0.02).

### TCR-seq analysis

*scIsoTyper* was used to assign most probable BCR IGH and IGK/L chains per droplet (based on nUMIs) and most probable TCR TRA and TRB chains per droplet (based on nUMIs)^57^. Briefly, *scIsoTyper* performs the following steps: (1) batches TCR data for IMGT submission; (2) parses the IMGT annotation results, for the quantification of V/J gene usages, and CDR3 sequence identity; (3) identifies the highest expressed alpha chain and beta chain TCR sequences per droplet; (3) clonality analysis; and (4) all this information is brought together in a meta-data format to include as part of the Seurat analysis, and provides statistics on the number of droplets with heavy and/or light chains and TCR alpha and/or beta chains.

### Differential gene expression analysis and pathway analysis

Differential gene expression analysis was performed using Seurat’s *FindMarkers()* function with the Poisson generalised linear model (GLM). We compared gene expression between two predefined cell clusters or conditions. The Poisson model was selected due to the discrete nature of the single-cell RNA-seq count data and its suitability for low-expression genes. Differentially expressed genes were defined as adjusted p-values <0.05.

The per cell pathway scores for each cell (which quantifies the feature expression programme for each pathway molecule) was calculated using the AddModuleScore using each pathway gene set. The statistics between the levels of each per-cell pathway score between samples was performed using two-sided MANOVA, shown to be statistically significant for each (p-values<1e-10). The mean pathway scores were then determined for each sample for plotting dynamics.

### Public datasets

Publicly available RNA sequencing datasets for SEM and RS4;11 cell lines were obtained from NCBI (GSE149158)^37^ and Cancer Cell Line Encyclopedia (CCLE) DepMap 2019^38^ (https://depmap.org/portal).

Transcripts per million (TPM) values for the following 14 genes were extracted from both datasets and collated: PROM1 (CD133), CD19, HLA-A, HLA-B, HLA-C, MICA, MICB, ULBP1, ULBP2, ULBP3, RAET1E (ULBP4), RAET1G (ULBP5) and RAET1L (ULBP6) and CD1D.

For NCBI (GSE149158), GEO2R was used to calculate the normalised TPM for each gene across triplicate samples for SEM (GSM4491229, GSM4491230 and GSM4491231) and RS4;11 (GSM4491211, GSM4491212 and GSM4491213) and the average TPM were subsequently enumerated. For CCLE DepMap, the TPM values for both cell lines were extracted from CCLE 2019 dataset “CCLE RNAseq gene expression data for 1019 cell lines (RSEM, gene)”.

### Data availability

Single cell RNA/TCR datasets have been deposited in Single Cell Portal under the accession: SCP2844, SCP2845, SCP2846, SCP2847, SCP2848 and are available via this link: (https://singlecell.broadinstitute.org/)

### Statistics

Two-tailed Mann-Whitney, Log-rank (Mantel-Cox) tests and ANOVA followed by multiple comparisons testing were used to compare experimental groups as indicated in the figure legends. Statistical analyses were performed using GraphPad Prism v10. Data are expressed as mean ± SD unless otherwise indicated.

## Acknowledgements

The work presented here was supported by a Cancer Research UK Children and Young People’s Cancer Innovation Award grant, co-funded by Children with Cancer (DRCPGM\100058) awarded to AR and AK with HR, NE, and BL also supported by the same grant. MUSS is supported by the Science and Technology Fellowship Trust, Government of Bangladesh. NEO and SI were supported by a donation from Azaylia Foundation.

TAM is supported by the Medical Research Council (MRC, UK) Molecular Haematology Unit grant MC_UU_00029/6. AR is supported by a Wellcome Trust Clinical Research Career Development Fellowship (216632/Z/19/Z) and Medical Research Council (MRC, UK) Molecular Haematology Unit grant MC_UU_00029/7. CH is supported by a Cancer Research UK Programme Foundation Award (DRCPFA-Nov21\100001).

The human fetal material was provided by the Joint MRC/Wellcome Trust Grant 099175/Z/ 12/Z Human Developmental Biology Resource (http://hdbr.org). ALL patient samples used in this study were provided by VIVO Biobank, supported by Cancer Research UK & Blood Cancer UK (Grant no. CRCPSC-Dec21\100003). We acknowledge contributions from the Imperial National Institute of Research Biomedical Research Centres. We are grateful for the technical support received from the WIMM Flow Cytometry Core, WIMM Single Cell facility, and John Radcliffe Biomedical Sciences Department.

The authors would like to thank Lynn Stevenson, University of Glasgow, for brain histology. The brain histology slides were scanned by the Glasgow Tissue Research Facility at the Queen Elizabeth University Hospital, Glasgow.

## Author contributions

HR, NE, BL, JWC and LF designed and performed experiments, analysed data, contributed to the preparation of the manuscript. KP, SR, TJ and IL performed experiments. MUSS and RBR analysed and visualised single cell data and contributed to the preparation of the manuscript. IAG contributed to the writing and editing of the manuscript. NEO, RT and SI performed immunophenotypic analysis of clinical samples. JB, OS and JB provided clinical samples, and contributed to the writing of the manuscript. CH supervised immunohistochemical analysis, interpreted data and contributed to the preparation of the manuscript. TM, AR and AK obtained funding, provided overall oversight of the experimental work, design, analysis, interpretation and wrote the manuscript.

---

**Suppl Table 3:**
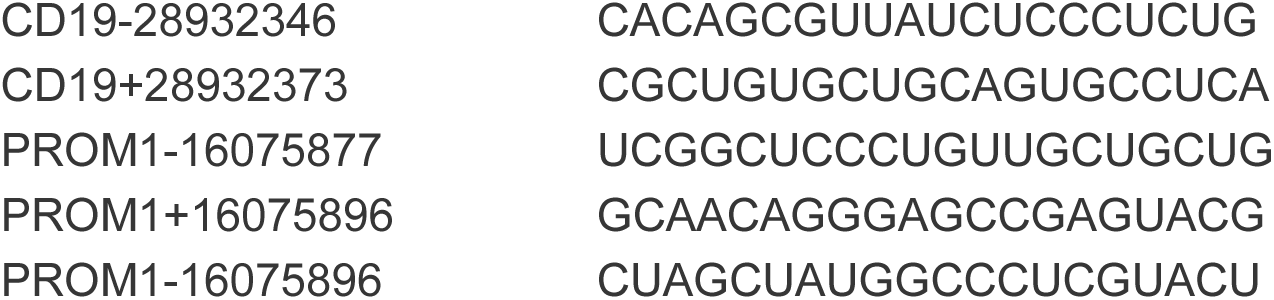
Guides for CRISPR knockout.

**Suppl_Table4.**
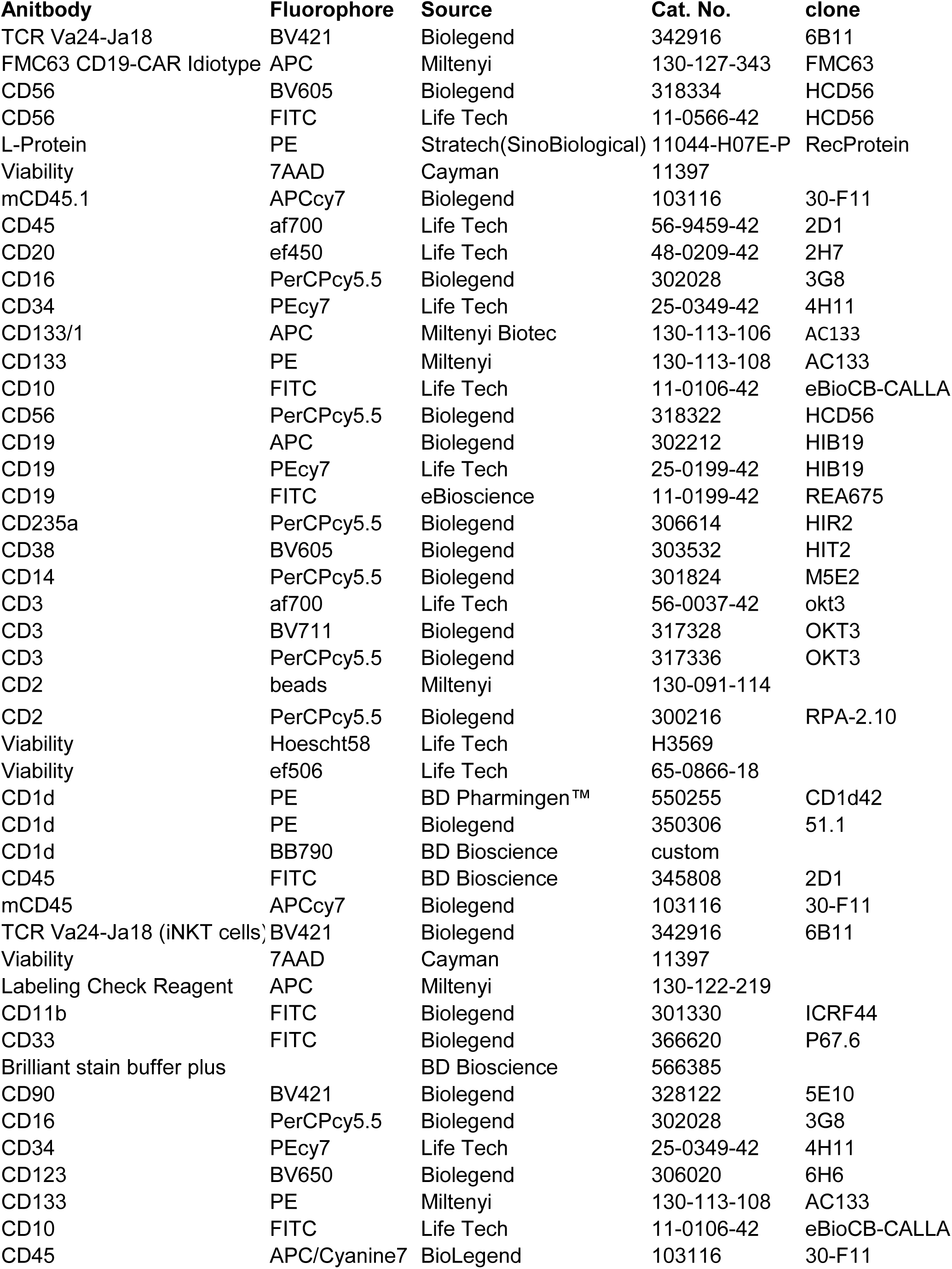
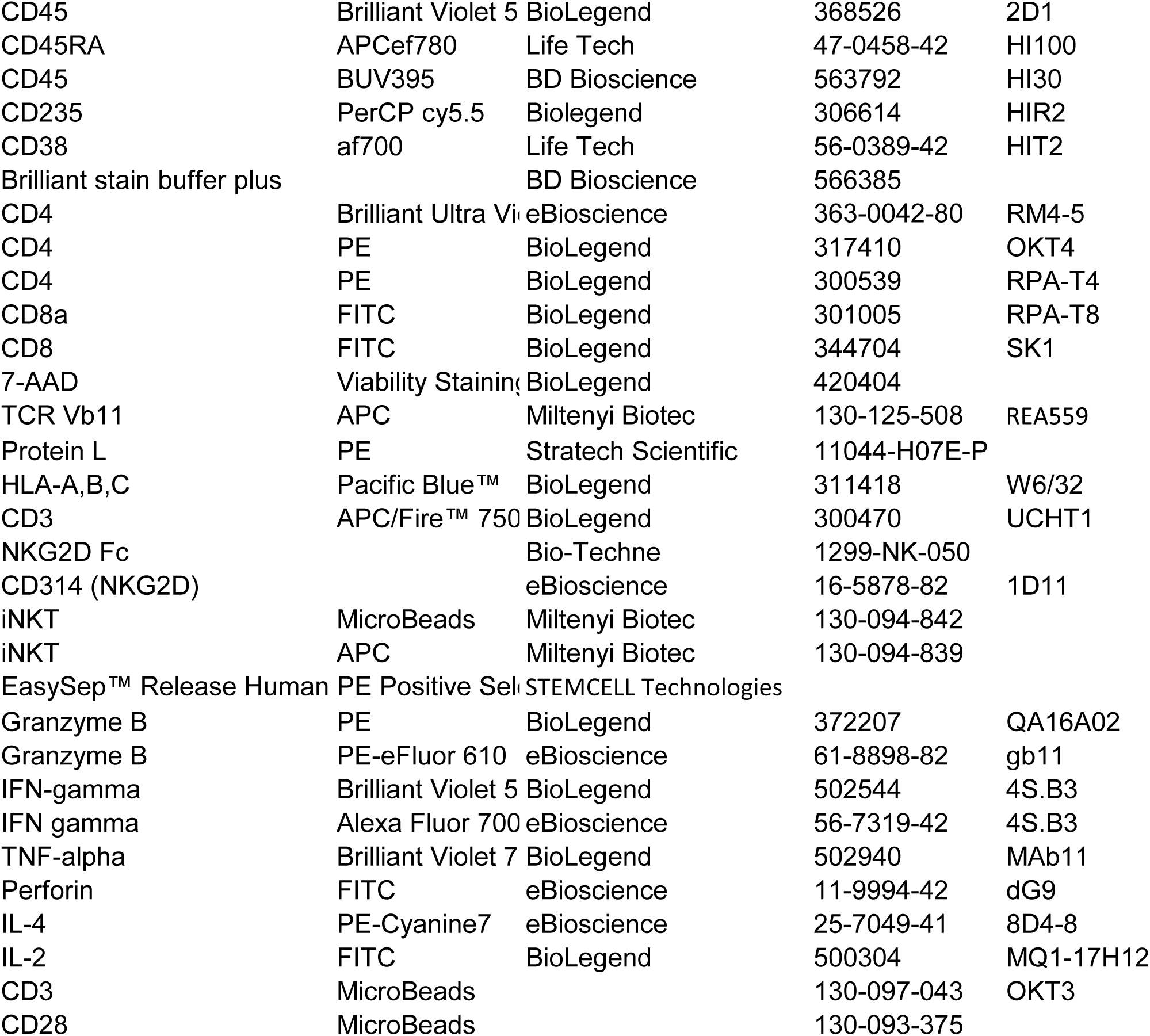
Antibody list.

| Anitbody | Fluorophore | Source | Cat. No. | clone |
| --- | --- | --- | --- | --- |
| TCR Va24-Ja18 | BV421 | Biolegend | 342916 | 6B11 |
| FMC63 CD19-CAR Idiotyp | APC | Miltenyi | 130-127-343 | FMC63 |
| CD56 | BV605 | Biolegend | 318334 | HCD56 |
| CD56 | FITC | Life Tech | 11-0566-42 | HCD56 |
| L-Protein | PE | Stratech(SinoBiological) | 11044-H07E-P | RecProtein |
| Viability | 7AAD | Cayman | 11397 |  |
| mCD45.1 | APCcy7 | Biolegend | 103116 | 30-F11 |
| CD45 | af700 | Life Tech | 56-9459-42 | 2D1 |
| CD20 | ef450 | Life Tech | 48-0209-42 | 2H7 |
| CD16 | PerCPcy5.5 | Biolegend | 302028 | 3G8 |
| CD34 | PEcy7 | Life Tech | 25-0349-42 | 4H11 |
| CD133/1 | APC | Miltenyi Biotec | 130-113-106 | AC133 |
| CD133 | PE | Miltenyi | 130-113-108 | AC133 |
| CD10 | FITC | Life Tech | 11-0106-42 | eBioCB-CALLA |
| CD56 | PerCPcy5.5 | Biolegend | 318322 | HCD56 |
| CD19 | APC | Biolegend | 302212 | HIB19 |
| CD19 | PEcy7 | Life Tech | 25-0199-42 | HIB19 |
| CD19 | FITC | eBioscience | 11-0199-42 | REA675 |
| CD235a | PerCPcy5.5 | Biolegend | 306614 | HIR2 |
| CD38 | BV605 | Biolegend | 303532 | HIT2 |
| CD14 | PerCPcy5.5 | Biolegend | 301824 | M5E2 |
| CD3 | af700 | Life Tech | 56-0037-42 | okt3 |
| CD3 | BV711 | Biolegend | 317328 | OKT3 |
| CD3 | PerCPcy5.5 | Biolegend | 317336 | OKT3 |
| CD2 | beads | Miltenyi | 130-091-114 |  |
| CD2 | PerCPcy5.5 | Biolegend | 300216 | RPA-2.10 |
| Viability | Hoescht58 | Life Tech | H3569 |  |
| Viability | ef506 | Life Tech | 65-0866-18 |  |
| CD1d | PE | BD Pharmingen™ | 550255 | CD1d42 |
| CD1d | PE | Biolegend | 350306 | 51.1 |
| CD1d | BB790 | BD Bioscience | custom |  |
| CD45 | FITC | BD Bioscience | 345808 | 2D1 |
| mCD45 | APCcy7 | Biolegend | 103116 | 30-F11 |
| TCR Va24-Ja18 (iNKT cells) | BV421 | Biolegend | 342916 | 6B11 |
| Viability | 7AAD | Cayman | 11397 |  |
| Labeling Check Reagent | APC | Miltenyi | 130-122-219 |  |
| CD11b | FITC | Biolegend | 301330 | ICRF44 |
| CD33 | FITC | Biolegend | 366620 | P67.6 |
| Brilliant stain buffer plus |  | BD Bioscience | 566385 |  |
| CD90 | BV421 | Biolegend | 328122 | 5E10 |
| CD16 | PerCPcy5.5 | Biolegend | 302028 | 3G8 |
| CD34 | PEcy7 | Life Tech | 25-0349-42 | 4H11 |
| CD123 | BV650 | Biolegend | 306020 | 6H6 |
| CD133 | PE | Miltenyi | 130-113-108 | AC133 |
| CD10 | FITC | Life Tech | 11-0106-42 | eBioCB-CALLA |
| CD45 | APC/Cyanine7 | BioLegend | 103116 | 30-F11 |
| CD45 | Brilliant Violet 5 | BioLegend | 368526 | 2D1 |
| CD45RA | APCef780 | Life Tech | 47-0458-42 | HI100 |
| CD45 | BUV395 | BD Bioscience | 563792 | HI30 |
| CD235 | PerCP cy5.5 | Biolegend | 306614 | HIR2 |
| CD38 | af700 | Life Tech | 56-0389-42 | HIT2 |
| Brilliant stain buffer plus |  | BD Bioscience | 566385 |  |
| CD4 | Brilliant Ultra Vi | eBioscience | 363-0042-80 | RM4-5 |
| CD4 | PE | BioLegend | 317410 | OKT4 |
| CD4 | PE | BioLegend | 300539 | RPA-T4 |
| CD8a | FITC | BioLegend | 301005 | RPA-T8 |
| CD8 | FITC | BioLegend | 344704 | SK1 |
| 7-AAD | Viability Staining | BioLegend | 420404 |  |
| TCR Vb11 | APC | Miltenyi Biotec | 130-125-508 | REA559 |
| Protein L | PE | Stratech Scientific | 11044-H07E-P |  |
| HLA-A,B,C | Pacific Blue™ | BioLegend | 311418 | W6/32 |
| CD3 | APC/Fire™ 750 | BioLegend | 300470 | UCHT1 |
| NKG2D Fc |  | Bio-Techne | 1299-NK-050 |  |
| CD314 (NKG2D) |  | eBioscience | 16-5878-82 | 1D11 |
| iNKT | MicroBeads | Miltenyi Biotec | 130-094-842 |  |
| iNKT | APC | Miltenyi Biotec | 130-094-839 |  |
| EasySep™ Release Human | PE Positive Sel | STEMCELL Technologies |  |  |
| Granzyme B | PE | BioLegend | 372207 | QA16A02 |
| Granzyme B | PE-eFluor 610 | eBioscience | 61-8898-82 | gb11 |
| IFN-gamma | Brilliant Violet 5 | BioLegend | 502544 | 4S.B3 |
| IFN gamma | Alexa Fluor 700 | eBioscience | 56-7319-42 | 4S.B3 |
| TNF-alpha | Brilliant Violet 7 | BioLegend | 502940 | MAb11 |
| Perforin | FITC | eBioscience | 11-9994-42 | dG9 |
| IL-4 | PE-Cyanine7 | eBioscience | 25-7049-41 | 8D4-8 |
| IL-2 | FITC | BioLegend | 500304 | MQ1-17H12 |
| CD3 | MicroBeads |  | 130-097-043 | OKT3 |
| CD28 | MicroBeads |  | 130-093-375 |  |

