## supplemental figs for "Bi-specific CAR-iNKT cell immunotherapy for high-risk KMT2A-rearranged leukemia outperforms CAR-T in an NKG2D-dependent manner and eradicates leptomeningeal disease"

[1]Centre for Haematology, Department of Immunology and Inflammation, Imperial College London, London, UK
[2]Department of Paediatrics, University of Oxford, Oxford, UK
[3]Department of Biochemistry, University of Oxford, Oxford, UK
[4]MRC Molecular Haematology Unit, MRC Weatherall Institute of Molecular Medicine, University of Oxford, Oxford, UK
[5]Department of Haematology, Great Ormond Street Hospital, London, United Kingdom
[6]Trinity College, University of Dublin, Dublin, Ireland
[7]Systems Biology Ireland, School of Medicine, University College Dublin, Ireland
[8]Children's Health Ireland at Crumlin, Dublin, Ireland
[9] Wolfson Wohl Cancer Research Centre, School of Cancer Sciences, College of Medical Veterinary and Life Sciences, University of Glasgow, Glasgow, UK
[10]Cancer Research UK Oxford Centre, University of Oxford, Oxford, UK
[11]Department of Haematology, Imperial College Health NHS Trust, Hammersmith Hospital, London, UK
*equal contribution

**Correspondence**
Anastasios Karadimitris:
Anindita Roy:
Thomas Milne:

### Supplementary Figure Legends

**Suppl Figure 1 related to Fig 1 a.** Representative flow-cytometry plots showing variable CD133 and CD1d co-expression in CD19+ blasts from 3 primary KMT2Ar ALL patient samples (chALL; childhood ALL, iALL: infant ALL) and a human fetal liver (FL) HSPC derived $^{CRISPR}$KMT2A-AFF1 ALL model. **b.** Percentage of CD19+ blasts that express CD133 in KMT2Ar iALL (n=10), KMT2Ar chALL (n=5), and $^{CRISPR}$KMT2A-AF4 ALL (n=10). **c.** Percentage of CD19+ blasts that express CD1d in KMT2Ar iALL (n=4), KMT2Ar chALL (n=5), and $^{CRISPR}$KMT2A-AFF1 ALL (n=4). **d.** Data shown as percentage of primitive CD34+38- progenitors and CD34+CD19+ B-progenitors that express CD133 from fetal liver (FL, n=8) and fetal bone marrow (FBM, n=8).

**Suppl Figure 2 related to Fig 1 & 2. a.** Growth curve of CD19, CD133 and CD19-CD133 CAR-iNKT. **b.** Cytotoxicity of mono- and bi-specific CAR-iNKT against the CD19+CD133+ KMT2Ar RS4;11 leukemia cell line (left). Data are from two independent experiments using two different iNKT donors. . **c**. *In vivo* activity of the mono-specific CD133 CAR-iNKT in NSG mice engrafted with Luc-dsRED SEM cells. 5x$10^6$ CAR-iNKT were iv transferred to NSG mice or not on day 6 post-leukemia cell injection. Leukemia burden as assessed by BLI is shown. **d.** Immunophenotypic analysis of an SEM subline propagated *in vivo* in treatment-free mice. Variable co-expression of CD19 and CD133 in dsRed+ SEM cells. **e & f.** Leukemia burden and survival in mice first injected with the SEM subline shown in c followed by treatment with indicated numbers of bi-specific CD19-CD133 CAR-iNKT (n=7)**. g.** Representative flow-cytometric analysis of BM and spleen in sacrificed mice treated with bi-specific CAR-iNKT as shown in d-f.

**Suppl Figure 3 related to Fig 2. a**. Schematic of experiment shown b&c. **b.** Overall survival of mice treated as described in the schematic (n=5 mice per group). *p<0.05 **p<0.01 **c**. Frequency of SEM cells with the indicated phenotype in the BM, spleen and liver of mice engrafted with parental SEM cells co-expressing CD19 and CD133 admixed with CD19-CD133+ and CD19+CD133- SEM cells at an 8:1:1 ratio respectively and treated with $10^6$ mono- or bi-specific CAR-iNKT cells (n=5-7 mice per group). With reference to untreated controls, frequency of CD19-CD133+ and CD19+CD133- SEM cells is not significantly different amongst the mono-specific CAR-iNKT-treated animals; with the only exception being a significant higher fraction of CD19-CD133+ cells in the spleens of animals treated with CD19 CAR-iNKT (p=0.03 vs untreated control; one-way ANOVA).

**Suppl Figure 4 related to Fig 2. a. F**low-cytometric identification of SEM cells after staining with anti-HLA-ABC mAb. Right: cumulative data of leukemia burden in BM on the indicated timepoints (n=7 mice per group). **b. Flow-cytometric** examples for SEM (top) and iNKT (bottom) from both d6 and d12 high dose CAR-iNKT treated animals culled at day 60 and 50 respectively. **c&d.** Leukemia burden assessed by BLI and overall survival of mice treated with bi-specific CAR-iNKT on day 16 after leukemia transfer (n=4 mice per group). **e.** Representative immunophenotypic analysis and cumulative data of leukemia burden in BM and spleen in CAR-iNKT-treated and untreated mice as shown in c&d. Leukemia cells are identified as HLA-class I+ cells. **f.** Heads of two bi-specific CAR-iNKT-treated animals showing stromal thickening in meninges (arrows), top panel H&E, bottom panel anti human CD19, scale bar 100μm.

**Suppl Fig 5 related to Fig 3&4. a.** bi-specific CAR transduction of T and iNKT cells from the same donor. **b.** Cytotoxic activity at 4 and 24hrs of untransduced T and iNKT and of their bi-specific CAR-transduced counterparts against RS4;11 leukemia cells. **c.** Representative flow-cytometric analysis of intracellular cytokine production by CAR-iNKT and CAR-T after their 4 and 24hr co-culture with SEM cells. **d.** CD19 and CD133 mono-specific CAR-transduced and untransduced T and iNKT against the parental CD19+CD133+ SEM and RS4;11 cells and the CD19+CD133- KOPN8 KMT2Ar cells. **e.** CD1d surface expression as assessed by flow-cytometry in SEM and RS4;11 cells. The Jurkat T cell line that expresses CD1d is shown as positive control. **f**. 4 and 24hr cytotoxic activity of bi-specific CAR-T and CAR-iNKT against SEM and $^{CRISPR}$KMT2A-AFF1 cells. **g. F**requency of HLA-ABC+ cells in the peripheral blood of mice injected with $^{CRISPR}$KMT2A-AFF1 cells as assessed by flow-cytometry on day 6, before they were treated with CAR-T and CAR-iNKT as shown. Staining of peripheral blood of a mouse that was not injected with $^{CRISPR}$KMT2A-AFF1 is also shown. **h.** Cytotoxicity of indicated effectors against the PDX KMT2A-AFF1 ALL cells. **i.** Representative flow-cytometric example of day 14 blood PDX cells identified as CD19+CD133+ cells in mice that were either untreated or treated with CAR-iNKT subsequently (at day 15). a-i: Representative of two independent experiments.

**Suppl Figure 6 related to Fig 5. a.** mRNA expression of NKG2D ligands in SEM and RS4;11 cells as assessed by RNA-seq**. b.** 24hr cytotoxicity of bi-specific CAR-T and-iNKT that had been pre-cultured with SEM cells against SEM cells in the presence of 10mg of NKG2D-Fc protein or PBS control. **c.** Representative examples of the FACS analysis of data shown in Fig 4h. **e.** Representative flow-cytometric examples of NKG2D expression on CAR-iNKT/T pre-cultured with parental and gene-edited SEM cells as shown in Fig 4f&h. **d.** Upregulation of NKG2D in CAR-T vs CAR-iNKT after 24hr co-culture with $^{CRISPR}$KMT2A-AFF1 leukemia cell. **e.** Left: Bi-specific CAR-iNKT pre-cultured with $^{CRISPR}$KMT2A-AFF1 leukemia cells are subsequently more cytotoxic than CAR-T against **C**AR target-negative CD19-CD133-SEM cells in an NKG2D-dependent manner. Right: Cytotoxicity of CAR-iNKT/T that had not been pre-cultured with $^{CRISPR}$KMT2A-AFF1. Representative of two independent experiments using two different iNKT donors. **f.** mRNA expression of indicated genes as assessed by RNA-seq of paired presentation KMT2Ar lymphoblasts and relapse myeloid blasts.

**Suppl Figure 7 related to Fig 6. a.** Numbers of iNKT and SEM cells in the bone marrow of mice receiving bi-specific CAR-iNKT only or SEM leukemia cells and CAR-iNKT. **b.** Table showing the number of cells in which TCR transcripts were identified. **c.** Overlay of TCR (TRA)- expressing cells on the UMAP map. **d.** Clone size of the invariant TCRVa24Ja18 clonotype. **e.** GSEA of genes over-expressed in (top) clusters 0 & 1 and (bottom) between cells isolated from the bone marrow of leukemia-bearing- vs leukemia-free mice on day 15, with reference to MSigDB Hallmarks genesets. **f.** Relative expression of indicated genes in the different experimental subgroups**. g.** Volcano plot showing differential gene expression between CAR-iNKT cells isolated from the bone marrow of leukemia-bearing- vs leukemia-free mice on day 15 (log2FC>1 and padj< 0.05). **h.** Frequency of S phase in CAR-iNKT cells from leukemia-bearing and -free mice on day 3 and day 15. Fisher exact test. **I.** Projection of cell cycle related signatures on UMAP. Cluster 2 is the most enriched for proliferative cells (that cluster 2 is more proliferative than clusters 0 and 1. P-value= 2.894e-18, Fisher exact test).

**Suppl Figure 8 related to Fig 6. a.** Left: Flow-cytometric identification of iNKT cells in the BM at sacrifice of animals described in Fig 2a&b. Right: cumulative data for a. **b.** Schematic of expansion and functional analysis of iNKT cells form a. **c.** Purity and CAR expression by iNKT of day 9 post ex vivo selection and expansion. **d.** Cytotoxic activity at 4 and 24hr of day 16 ex vivo expanded CAR-iNKT against parental SEM, SEM with variable co-expression of CD19 and CD133 (BM; Suppl Fig 2d) and gene-edited SEM cells lacking expression of CD19 and CD133. **e.** 4 and 24hr cytotoxicity assay with day 23 CAR-iNKT against the CD19+CD133- C1R and C1R-CD1d cells in the presence or not of aGalCer (100ng/ml). Representative of two independent experiments.

**Suppl Figure 9. Impact of bi-specific CAR-iNKT on hematopoiesis in humanized mice**
**a.** Schematic of xenograft experiment to test hematological toxicity of CD19/CD133 CAR-iNK-T *in vivo.* **b.** Peripheral blood human CD45 engraftment levels in mice treated with PBS (n=5) or $10^7$ CD19/CD133 CAR-iNKT (n=7). **c)** Peripheral blood human CD19+ cells in mice treated with PBS (n=5) or $10^7$ CD19/CD133 CAR-iNKT (n=7), in two separate experiments, showing a transient drop in B cell proportions 1-3 days post CAR injection Unpaired t-test; *p<0.05. **d.** Long term engraftment in the bone marrow of mice treated with PBS or $10^7$ CD19/CD133 CAR-iNKT: from left to right: proportion of hCD45 cells in BM (PBS, n=4, CAR, n=6); B cells, T cells and myeloid cells in the BM expressed as proportion of hCD45+ cells (PBS, n=4, CAR, n=6); immature CD34+ cells in the BM expressed as proportion of hCD45+ cells (PBS, n=4, CAR, n=5).

**Suppl Fig 10. Analysis of peripheral blood and bone marrow of humanized mice receiving bi-specific CAR-iNKT. a.** representative flow plots showing gating strategy used to determine peripheral blood (PB) and bone marrow (BM) engraftment and lineage output. The data shown is from BM at cull for PBS treated (left) and CAR-iNKT treated (right) mice. **b.** representative flow plots showing frequency of CD19+ B cells and CD3+ T cells in the PB of control mice treated with PBS (top row) and mice treated with CAR-iNKT at 9 weeks (bottom row). Data shown as % of hCD45+ cells. **c.** representative flow plots showing gating strategy used to determine immature CD34+ cells in the BM at cull. **d.** proportion of hCD45 cells in BM of secondary engrafted NSG mice. Each mouse was injected with 1M whole BM from primary engrafted mice treated with PBS (n=4) or 10M CAR (n=4)

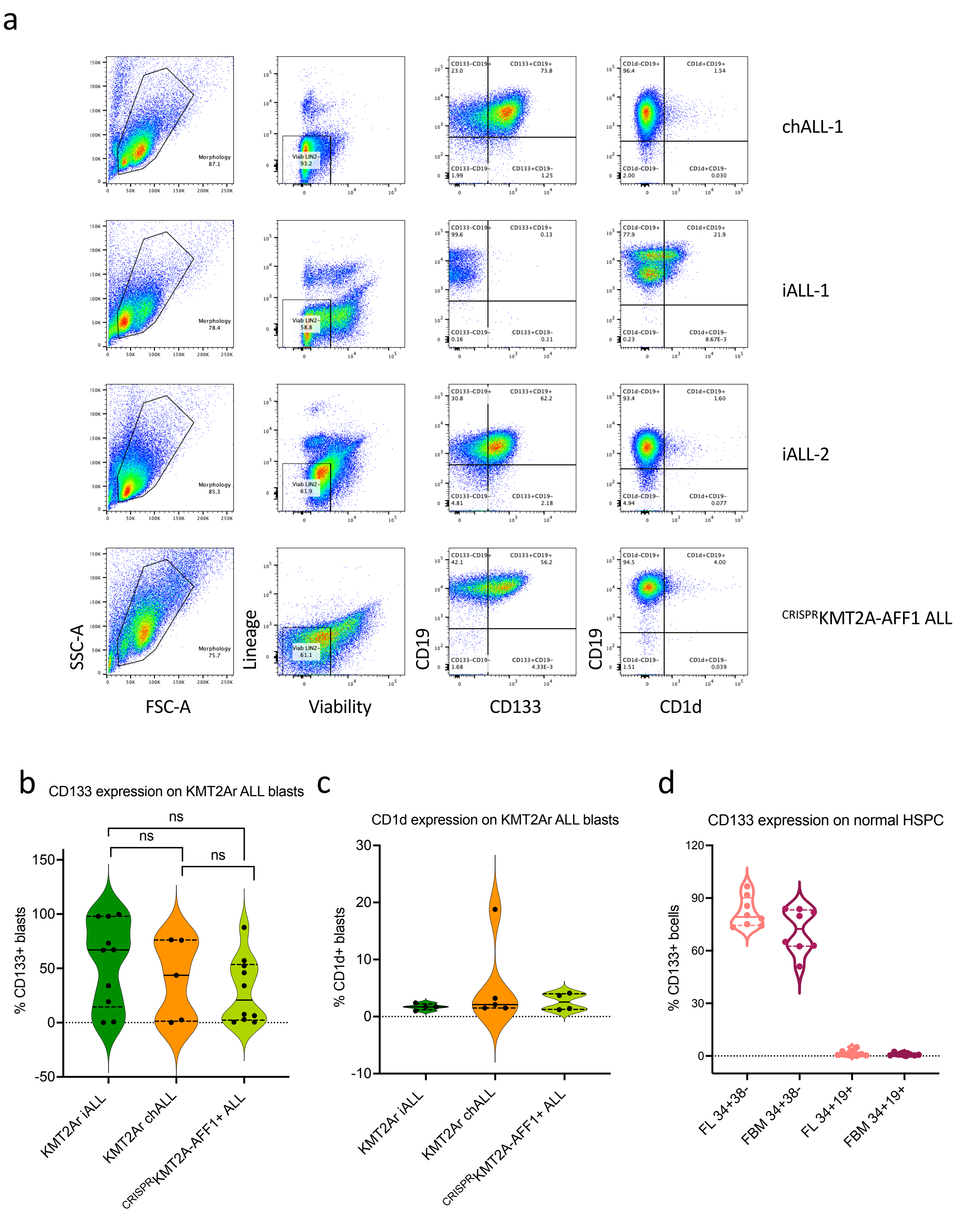


**Suppl Figure 1 related to Fig 1. a.** Representative flow-cytometry plots showing variable CD133 and CD1d co-expression in CD19+ blasts from 3 primary KMT2Ar ALL patient samples (chALL; childhood ALL, iALL: infant ALL) and a human fetal liver (FL) HSPC derived CRISPRKMT2A-AFF1 ALL model. **b.** Percentage of CD19+ blasts that express CD133 in KMT2Ar iALL (n=10), KMT2Ar chALL (n=5), and CRISPRKMT2A-AF4 ALL (n=10). **c.** Percentage of CD19+ blasts that express CD1d in KMT2Ar iALL (n=4), KMT2Ar chALL (n=5), and CRISPRKMT2A-AFF1 ALL (n=4). **d.** Data shown as percentage of primitive CD34+38- progenitors and CD34+CD19+ B-progenitors that express CD133 from fetal liver (FL, n=8) and fetal bone marrow (FBM, n=8).

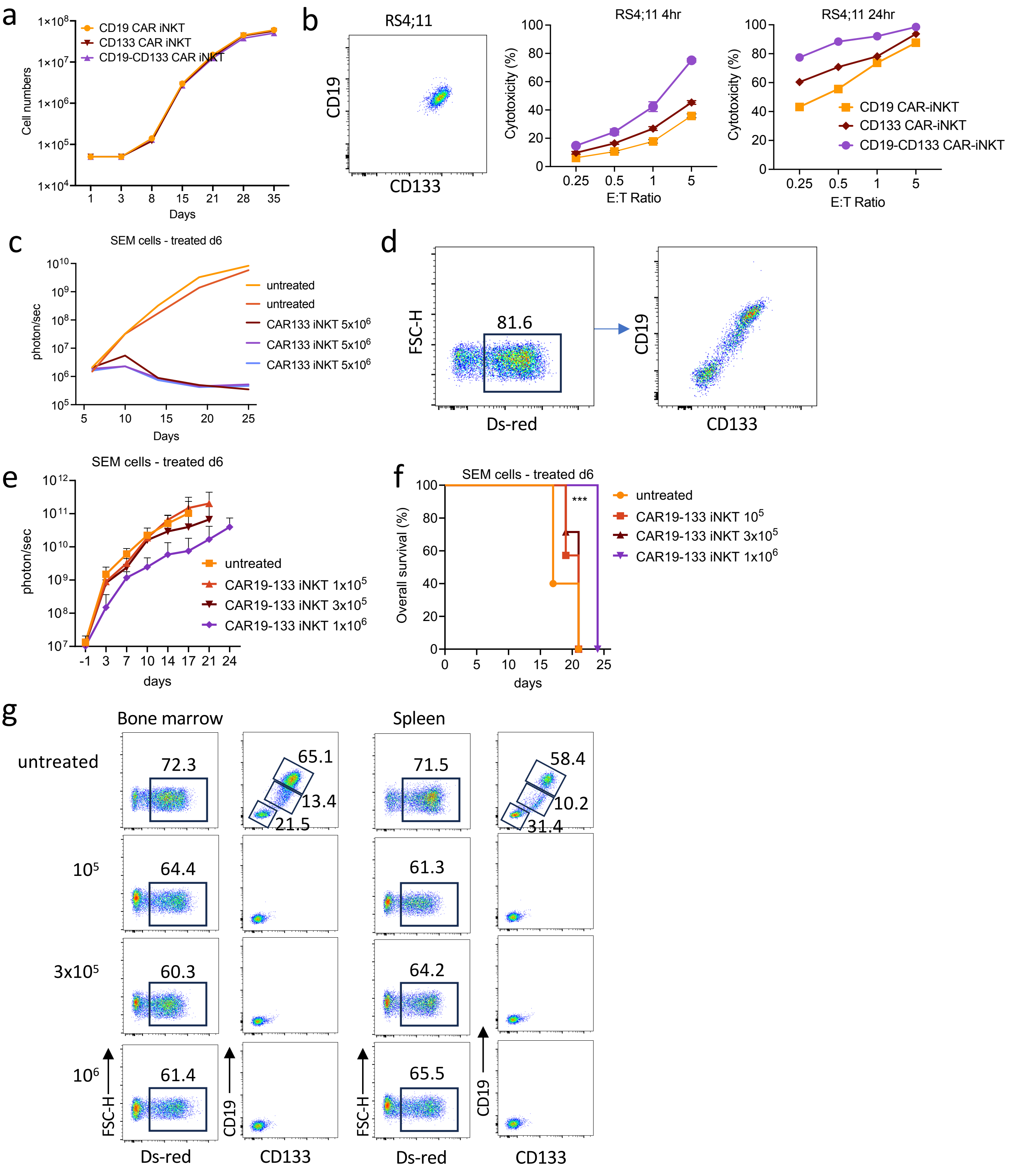


**Suppl Figure 2 related to Fig 1 & 2. a.** Growth curve of CD19, CD133 and CD19-CD133 CAR-iNKT. **b.** Cytotoxicity of mono- and bi-specific CAR-iNKT against the CD19+CD133+ KMT2Ar RS4;11 leukemia cell line (left). Data are from two independent experiments using two different iNKT donors. . **c**. In vivo activity of the mono-specific CD133 CAR-iNKT in NSG mice engrafted with Luc-dsRED SEM cells. $5x10^6$ CAR-iNKT were iv transferred to NSG mice or not on day 6 post-leukemia cell injection. Leukemia burden as assessed by BLI is shown. **d.** Immunophenotypic analysis of an SEM subline propagated in vivo in treatment-free mice. Variable co-expression of CD19 and CD133 in dsRed+ SEM cells. **e & f.** Leukemia burden and survival in mice first injected with the SEM subline shown in c followed by treatment with indicated numbers of bi-specific CD19-CD133 CAR-iNKT (n=7)**. g.** Representative flow-cytometric analysis of BM and spleen in sacrificed mice treated with bi-specific CAR-iNKT as shown in d-f.

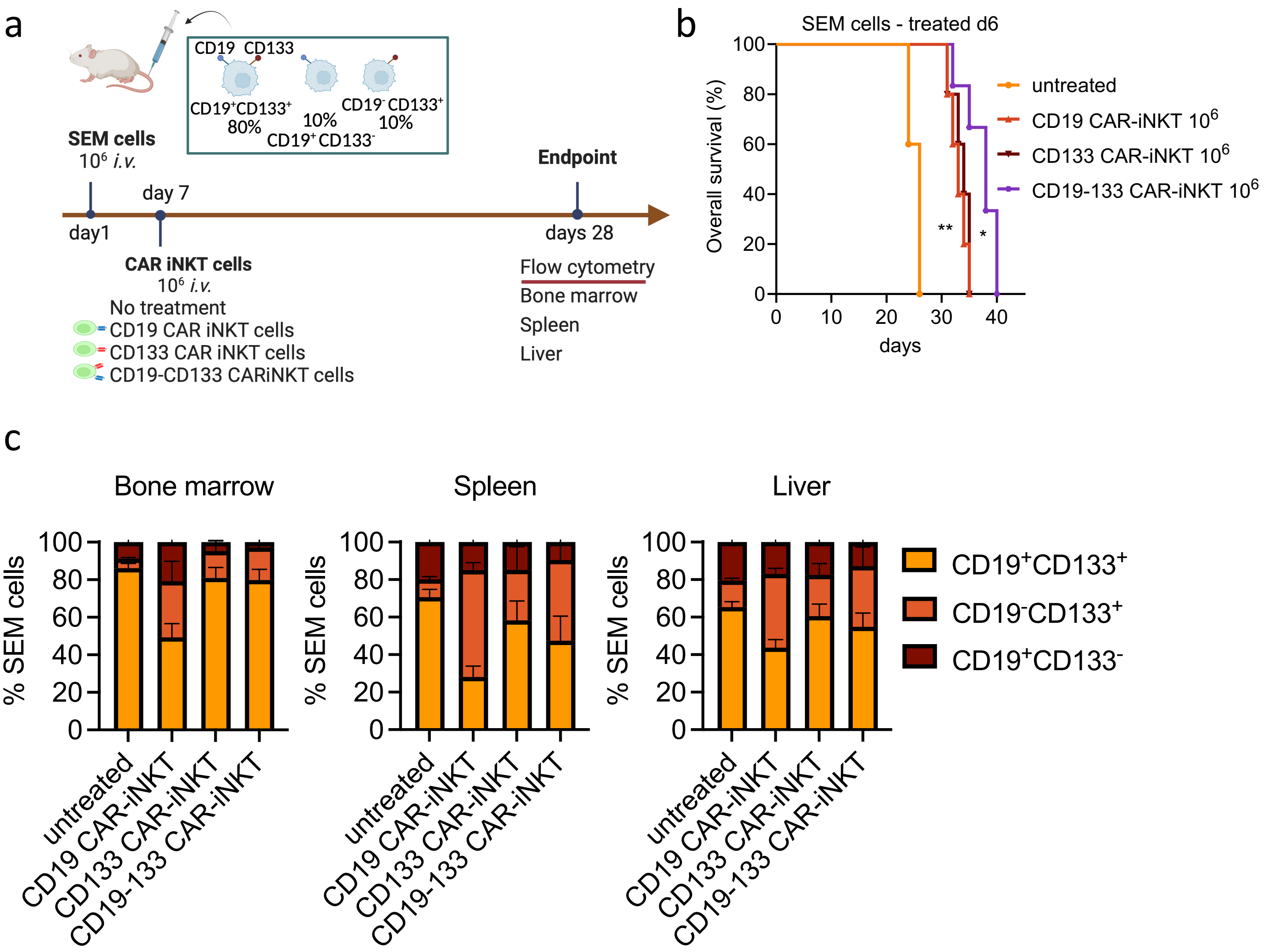


**Suppl Figure 3 related to Fig 2. a**. Schematic of experiment shown b&c. **b.** Overall survival of mice treated as described in the schematic (n=5 mice per group). *p<0.05 **p<0.01 **c**. Frequency of SEM cells with the indicated phenotype in the BM, spleen and liver of mice engrafted with parental SEM cells co-expressing CD19 and CD133 admixed with CD19-CD133+ and CD19+CD133- SEM cells at an 8:1:1 ratio respectively and treated with $10^6$ mono- or bi-specific CAR-iNKT cells (n=5-7 mice per group). With reference to untreated controls, frequency of CD19-CD133+ and CD19+CD133- SEM cells is not significantly different amongst the mono-specific CAR-iNKT-treated animals; with the only exception being a significant higher fraction of CD19-CD133+ cells in the spleens of animals treated with CD19 CAR-iNKT (p=0.03 vs untreated control; one-way ANOVA)

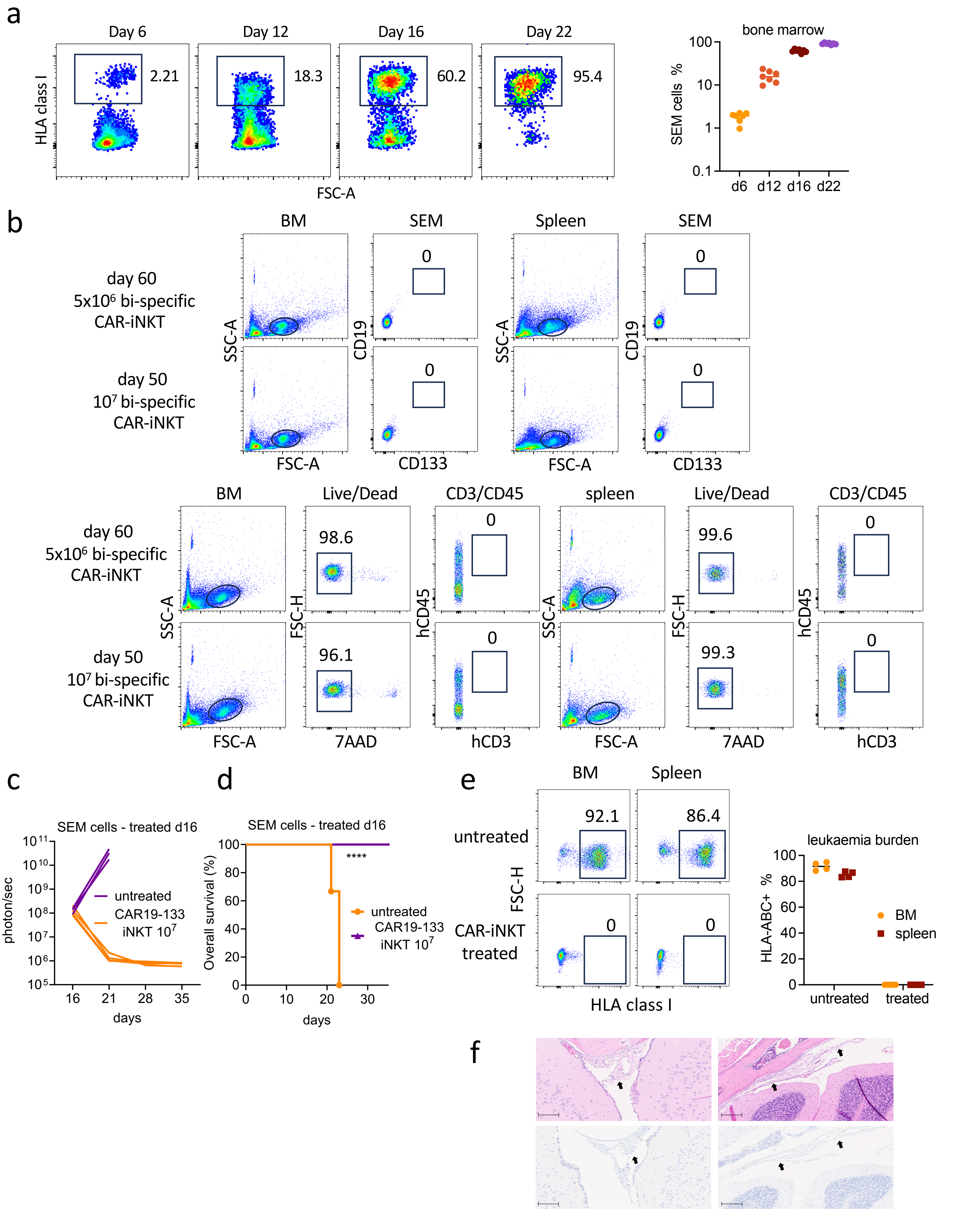


**Suppl Figure 4 related to Fig 2. a.** Flow-cytometric identification of SEM cells after staining with anti-HLA-ABC mAb. Right: cumulative data of leukemia burden in BM on the indicated timepoints (n=7 mice per group). **b.** Flow-cytometric examples for SEM (top) and iNKT (bottom) from both d6 and d12 high dose CAR-iNKT treated animals culled at day 60 and 50 respectively. **c&d.** leukemia burden assessed by BLI and overall survival of mice treated with bi-specific CAR-iNKT on day 16 after leukemia transfer (n=4 mice per group). **e.** Representative immunophenotypic analysis and cumulative data of leukemia burden in BM and spleen in CAR-iNKT-treated and untreated mice as shown in c&d. Leukemia cells are identified as HLA-class I+. **f.** Heads of two bi-specific CAR-iNKT-treated animals showing stromal thickening in meninges (arrows), top panel H&E, bottom panel anti human CD19, scale bar 100mm.

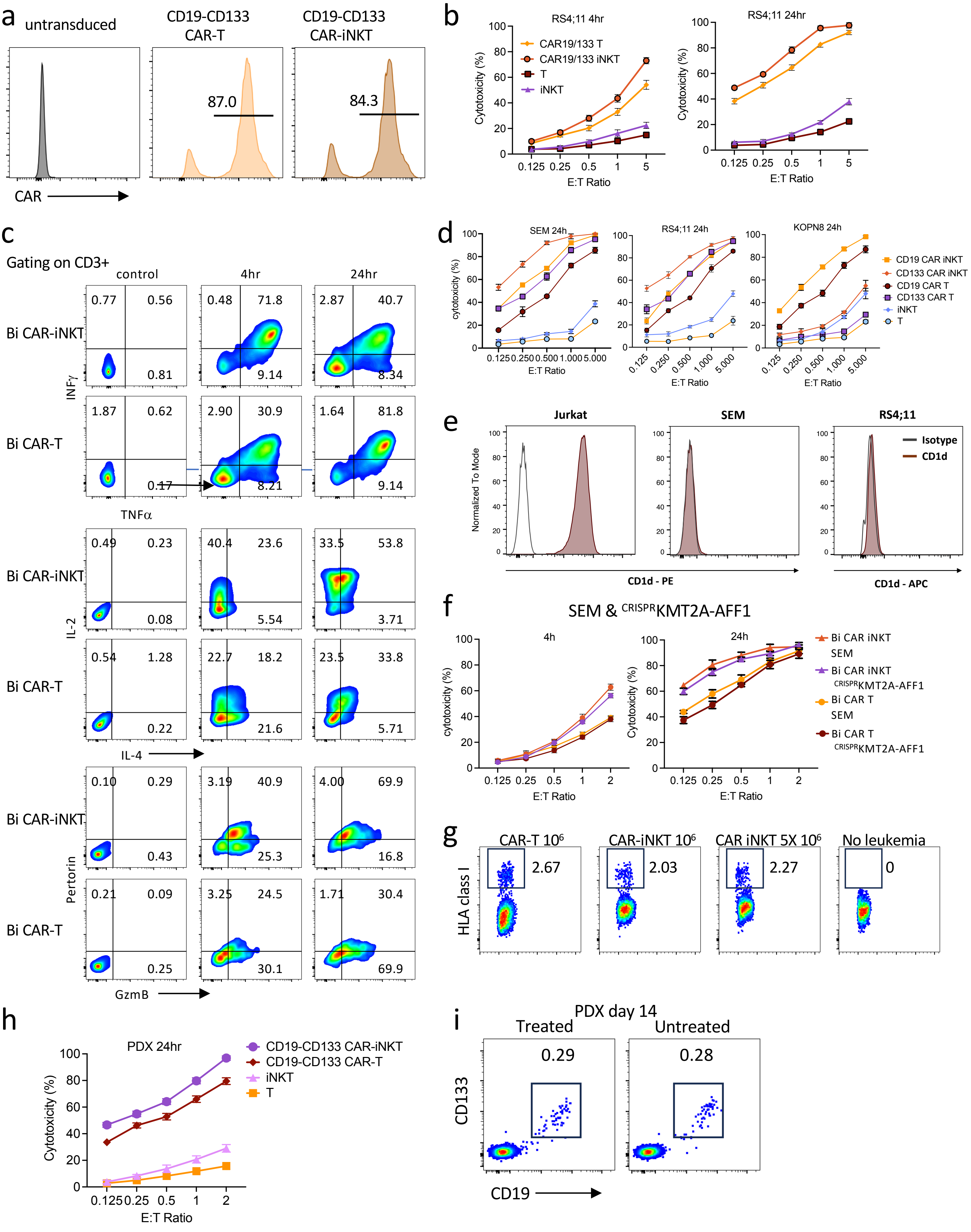


**Suppl Fig 5 related to Fig 3&4.**

**Suppl Fig 5 related to Fig 3&4. a.** bi-specific CAR transduction of T and iNKT cells from the same donor. **b.** Cytotoxic activity at 4 and 24hrs of untransduced T and iNKT and of their bi-specific CAR-transduced counterparts against RS4;11 leukemia cells. **c.** Representative flow-cytometric analysis of intracellular cytokine production by CAR-iNKT and CAR-T after their 4 and 24hr co-culture with SEM cells. **d.** CD19 and CD133 mono-specific CAR-transduced and untransduced T and iNKT against the parental CD19+CD133+ SEM and RS4;11 cells and the CD19+CD133- KOPN8 KMT2Ar cells. **e.** CD1d surface expression as assessed by flow-cytometry in SEM and RS4;11 cells. The Jurkat T cell line that expresses CD1d is shown as positive control. **f**. 4 and 24hr cytotoxic activity of bi-specific CAR-T and CAR-iNKT against SEM and $^{CRISPR}$KMT2A-AFF1 cells. **g. F**requency of HLA class I+ cells in the peripheral blood of mice injected with $^{CRISPR}$KMT2A-AFF1 cells as assessed by flow-cytometry on day 6, before they were treated with CAR-T and CAR-iNKT as shown. Staining of peripheral blood of a mouse that was not injected with $^{CRISPR}$KMT2A-AFF1 is also shown. **h.** Cytotoxicity of indicated effectors against the PDX KMT2A-AFF1 ALL cells. **i.** Representative flow-cytometric example of day 14 blood PDX cells identified as CD19+CD133+ cells in mice that were either untreated or treated with CAR-iNKT subsequently (at day 15). a-i: Representative of two independent experiments.

a

**SEM**

| Gene | TPM: GEO (GSE149158) | TPM: CCLE DepMap 2019 | |
|---|---|---|---|
| PROM1 (CD133) | 171.6 | 397.7 | SEM key signatures |
| CD19 | 146.4 | 219.2 | |
| HLA-A | 212.1 | 247.1 | MHC Class 1 |
| HLA-B | 406.2 | 448.2 | |
| HLA-C | 229.0 | 264.6 | |
| MICA | 8.7 | 11.6 | NKG2D ligands |
| MICB | 23.6 | 26.3 | |
| ULBP1 | 1.0 | 1.6 | |
| ULBP2 | 1.9 | 1.4 | |
| ULBP3 | 4.0 | 1.7 | |
| RAET1E (ULBP4) | 0.0 | 0.0 | |
| RAET1G (ULBP5) | 0.0 | 0.0 | |
| RAET1L (ULBP6) | 0.2 | 0.0 | |
| CD1D | 0.1 | 0.2 | |

**RS4;11**

| Gene | TPM: GEO (GSE149158) | TPM: CCLE DepMap 2019 | |
|---|---|---|---|
| PROM1 (CD133) | 124.5 | 136.0 | RS4;11 key signatures |
| CD19 | 84.9 | 133.3 | |
| HLA-A | 305.0 | 274.1 | MHC Class 1 |
| HLA-B | 534.3 | 471.7 | |
| HLA-C | 195.1 | 169.4 | |
| MICA | 7.4 | 7.0 | NKG2D ligands |
| MICB | 26.0 | 5.3 | |
| ULBP1 | 0.0 | 0.0 | |
| ULBP2 | 0.4 | 0.3 | |
| ULBP3 | 0.5 | 0.0 | |
| RAET1E (ULBP4) | 0.0 | 0.0 | |
| RAET1G (ULBP5) | 0.0 | 0.0 | |
| RAET1L (ULBP6) | 0.1 | 0.0 | |
| CD1D | 0.0 | 0.0 | |

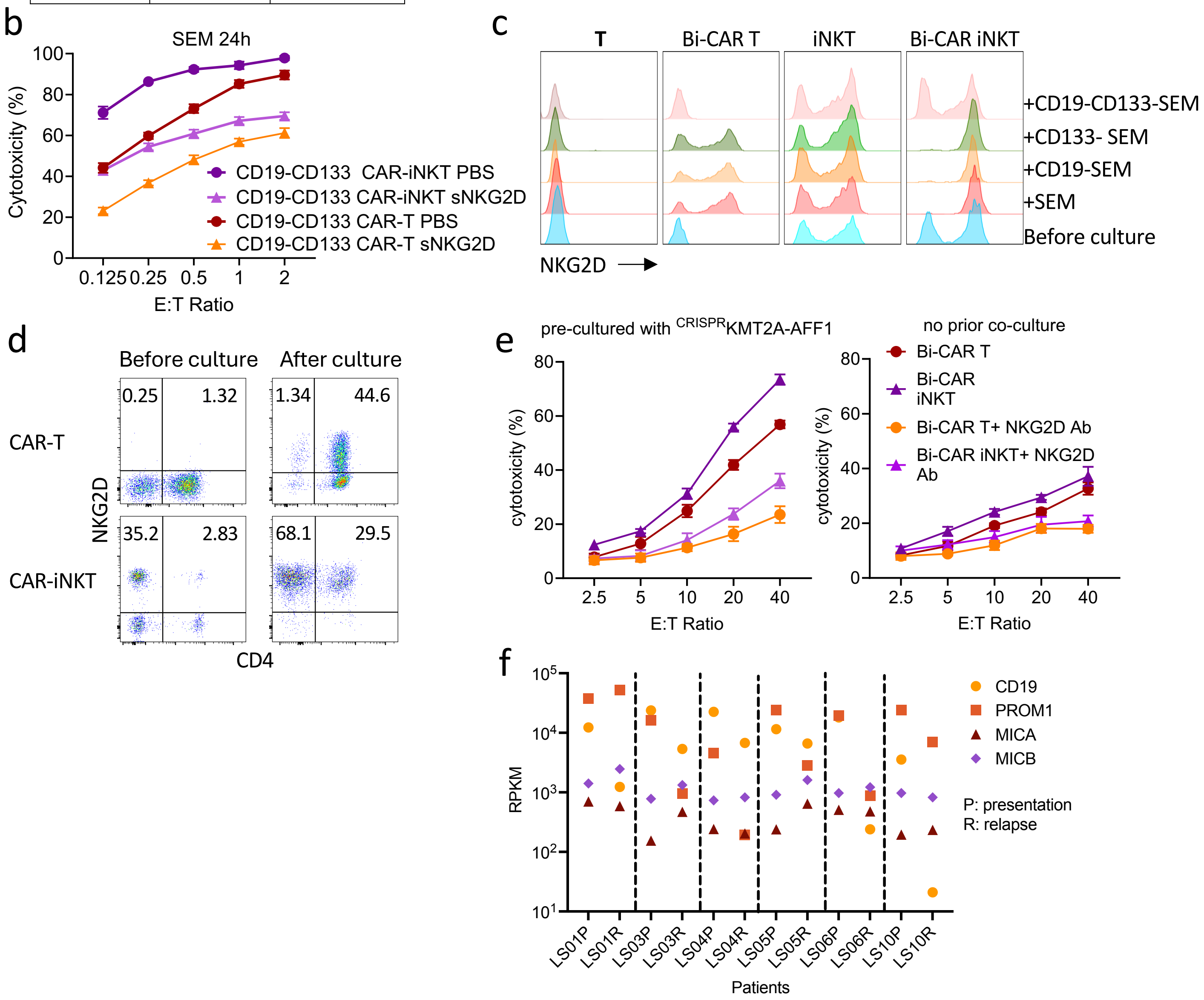


**Suppl Figure 6 related to Fig 5. a.** mRNA expression of NKG2D ligands in SEM and RS4;11 cells as assessed by RNA-seq. **b.** 24hr cytotoxicity of bi-specific CAR-T and-iNKT that had been pre-cultured with SEM cells against SEM cells in the presence of 10µg of NKG2D-Fc protein or PBS control. **c.** Representative examples of the FACS analysis of data shown in Fig 4h. **e.** Representative flow-cytometric examples of NKG2D expression on CAR-iNKT/T pre-cultured with parental and gene-edited SEM cells as shown in Fig 4f&h. **d.** Upregulation of NKG2D in CAR-T vs CAR-iNKT after 24hr co-culture with CRISPRKMT2A-AFF1 leukemia cell. **e.** Left: Bi-specific CAR-iNKT pre-cultured with CRISPRKMT2A-AFF1 leukemia cells are subsequently more cytotoxic than CAR-T against **C**AR target-negative CD19-CD133-SEM cells in an NKG2D-dependent manner. Right: Cytotoxicity of CAR-iNKT/T that had not been pre-cultured with CRISPRKMT2A-AFF1. Representative of two independent experiments using two different iNKT donors. **f.** mRNA expression of indicated genes as assessed by RNA-seq of paired presentation KMT2Ar lymphoblasts and relapse myeloid blasts.

a

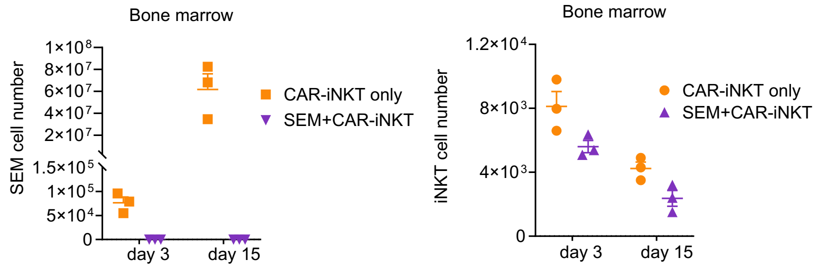


b

| | Total cells with GEX | TCR_status: TRA+ | TCR_status: TRA+TRB+ | TCR_status: TRB+ | TCR_status: none |
|---|---|---|---|---|---|
| BMd15_CAR_7489cells | 1526 | 94 | 472 | 138 | 822 |
| BMd15_CAR_SEM_2768 | 345 | 22 | 166 | 16 | 141 |
| BMd3_CAR_6349cells | 1115 | 55 | 565 | 126 | 369 |
| BMd3_CAR_SEM_7645 | 2141 | 124 | 964 | 275 | 778 |
| Naive_CAR_30100 | 8426 | 1244 | 5514 | 708 | 960 |

c

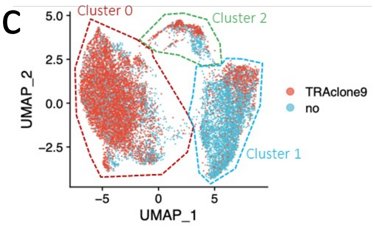


d

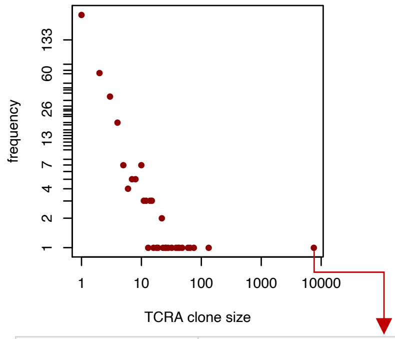


| | |
|---|---|
| TRA clone ID | Clone 9 |
| TRA V gene | TRAV10*01 |
| TRA J gene | TRAJ18*01 |
| CDR3 regions (amino acid) | CVVSDRGSTLGRLYF |
| CDR3 regions (nucleotide) | TGTGTGGTGAGCGACAGAGGCTCAACCCTGGGGAGGCTATACTTT |

e

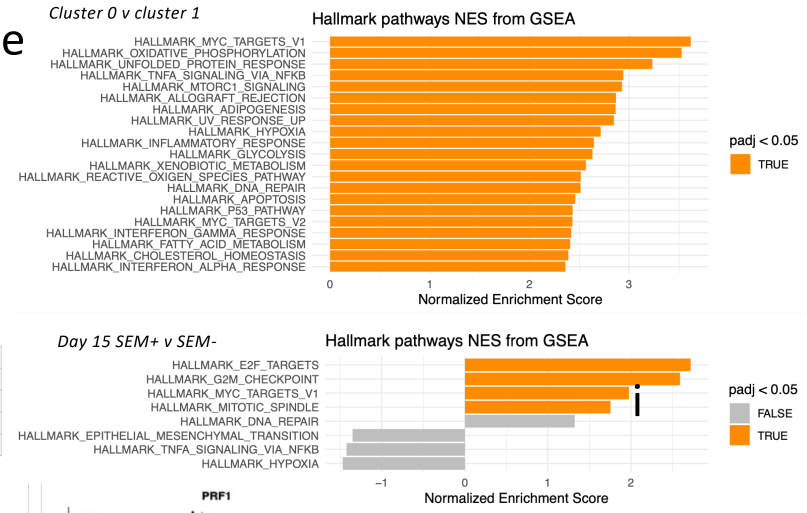


f

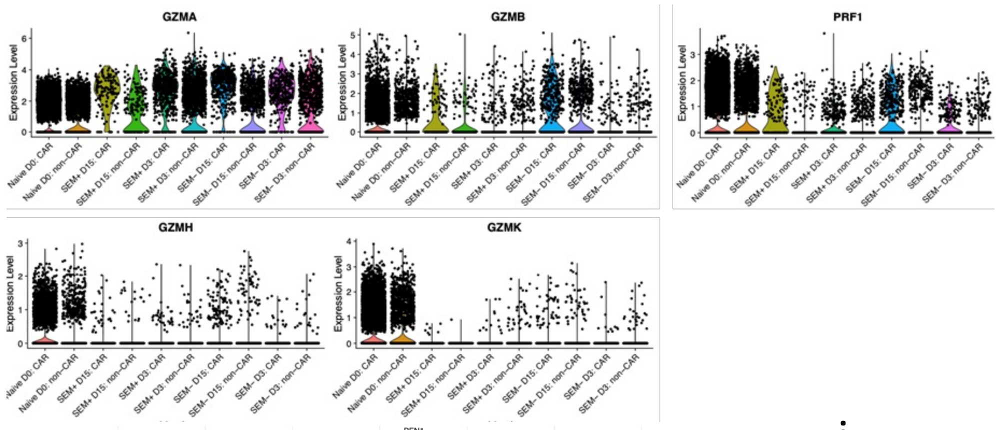


g

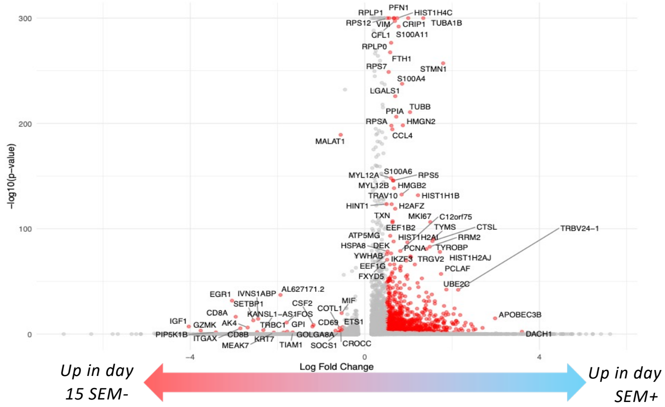


i

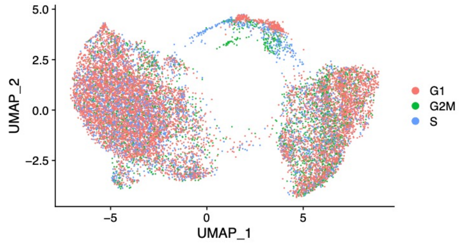


**Number of cells (%)**

| | Cluster 0 | Cluster 1 | Cluster 2 |
|---|---|---|---|
| Non-S phase | 6101 (75.4%) | 3677 (78.6%) | 486 (62.1%) |
| S phase | 1992 (24.6%) | 1001 (21.4%) | 296 (37.9%) |

h

| Sample | Number (%) of cells in non-S phase | Number (%) of cells in S phase | p-value (between SEM+/-) |
|---|---|---|---|
| BMd3_CAR_SEM_7645 | 1541 (72%) | 600 (28%) | 0.3268 |
| BMd3_CAR_6349cells | 784 (70.3%) | 331 (29.7%) | |
| BMd15_CAR_SEM_2768 | 195 (56.5%) | 150 (43.5%) | 6.20E-06 |
| BMd15_CAR_7489cells | 1060 (69.5%) | 466 (30.5%) | |

**Suppl Fig 7 related to Fig 6.**

**Suppl Figure 7 related to Fig 6. a.** Numbers of iNKT and SEM cells in the bone marrow of mice receiving bi-specific CAR-iNKT only or SEM leukemia cells and CAR-iNKT. **b.** Table showing the number of cells in which TCR transcripts were identified. **c.** Overlay of TCR (TRA)- expressing cells on the UMAP map. **d.** Clone size of the invariant TCRVa24Ja18 clonotype. **e.** GSEA of genes over-expressed in (top) clusters 0 & 1 and (bottom) between cells isolated from the bone marrow of leukemia-bearing- vs leukemia-free mice on day 15, with reference to MSigDB Hallmarks genesets. **f.** Relative expression of indicated genes in the different experimental subgroups**. g.** Volcano plot showing differential gene expression between CAR-iNKT cells isolated from the bone marrow of leukemia-bearing- vs leukemia-free mice on day 15 (log2FC>1 and padj< 0.05). **h.** Frequency of S phase in CAR-iNKT cells from leukemia-bearing and -free mice on day 3 and day 15. Fisher exact test. **I.** Projection of cell cycle related signatures on UMAP. Cluster 2 is the most enriched for proliferative cells (that cluster 2 is more proliferative than clusters 0 and 1. P-value= 2.894e-18, Fisher exact test).

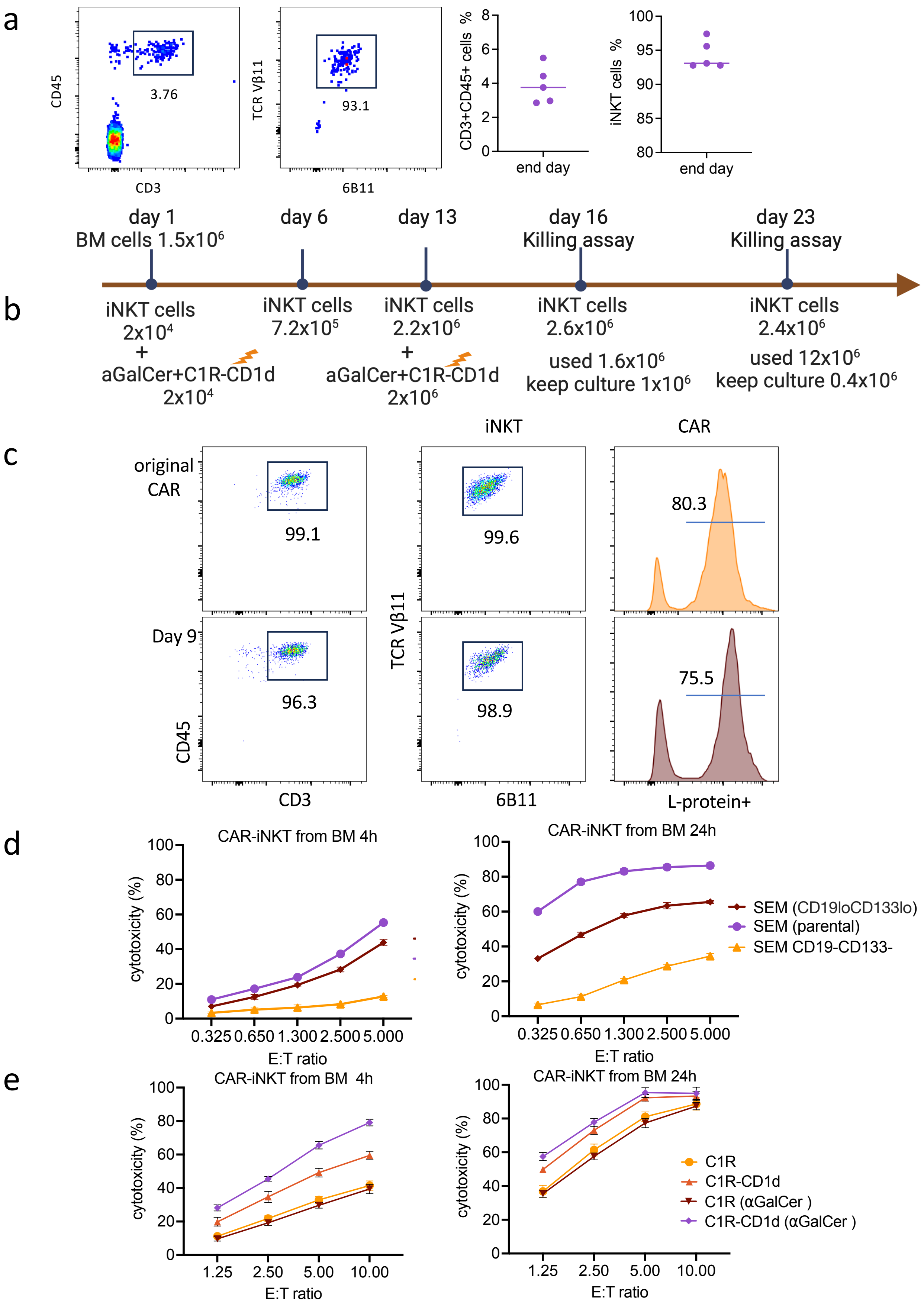


**Suppl Figure 8 related to Fig 6. a.** Left: Flow-cytometric identification of iNKT cells in the BM at sacrifice of animals described in Fig 2a&b. Right: cumulative data for a. **b.** Schematic of expansion and functional analysis of iNKT cells form a. **c.** Purity and CAR expression by iNKT of day 9 post ex vivo selection and expansion. **d.** Cytotoxic activity at 4 and 24hr of day 16 ex vivo expanded CAR-iNKT against parental SEM, SEM with variable co-expression of CD19 and CD133 (BM; Suppl Fig 2d) and gene-edited SEM cells lacking expression of CD19 and CD133. **e.** 4 and 24hr cytotoxicity assay with day 23 CAR-iNKT against the CD19+CD133- C1R and C1R-CD1d cells in the presence or not of aGalCer (100ng/ml). Representative of two independent experiments.

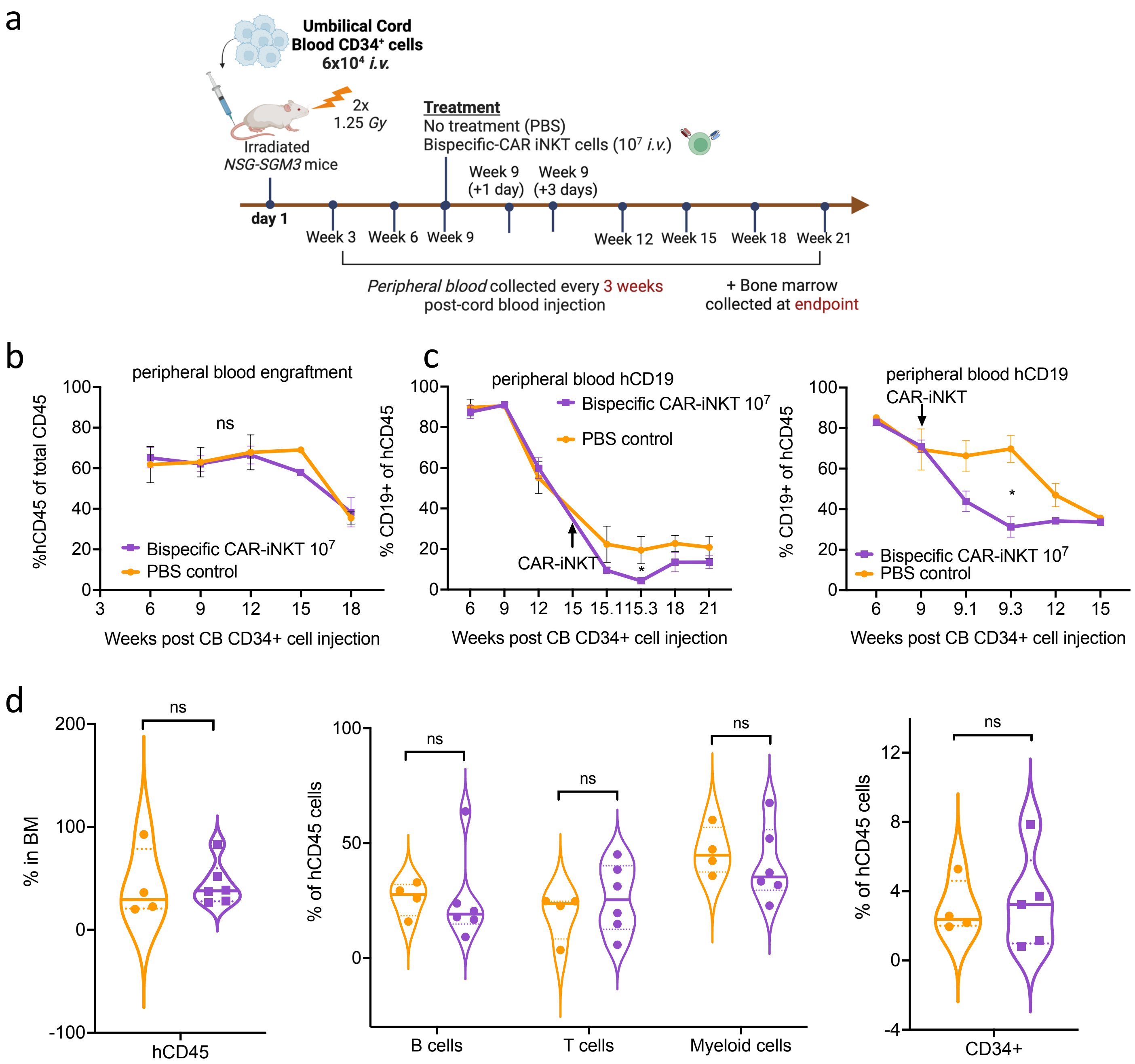


**Suppl Figure 9. Impact of bi-specific CAR-iNKT on hematopoiesis in humanized mice. a.** Schematic of xenograft experiment to test hematological toxicity of CD19-CD133 CAR-iNK-T *in vivo.* **B.** Peripheral blood human CD45 engraftment levels in mice treated with PBS (n=5) or $10^7$ CD19-CD133 CAR-iNKT (n=7). **c)** Peripheral blood human CD19+ cells in mice treated with PBS (n=5) or $10^7$ CD19-CD133 CAR-iNKT (n=7), in two separate experiments, showing a transient drop in B cell proportions 1-3 days post CAR injection Unpaired t-test; *$p<0.05$. **d.** Long term engraftment in the bone marrow of mice treated with PBS or $10^7$ CD19-CD133 CAR-iNKT: from left to right: proportion of hCD45 cells in BM (PBS, n=4, CAR, n=6); B cells, T cells and myeloid cells in the BM expressed as proportion of hCD45+ cells (PBS, n=4, CAR, n=6); immature CD34+ cells in the BM expressed as proportion of hCD45+ cells (PBS, n=4, CAR, n=5)

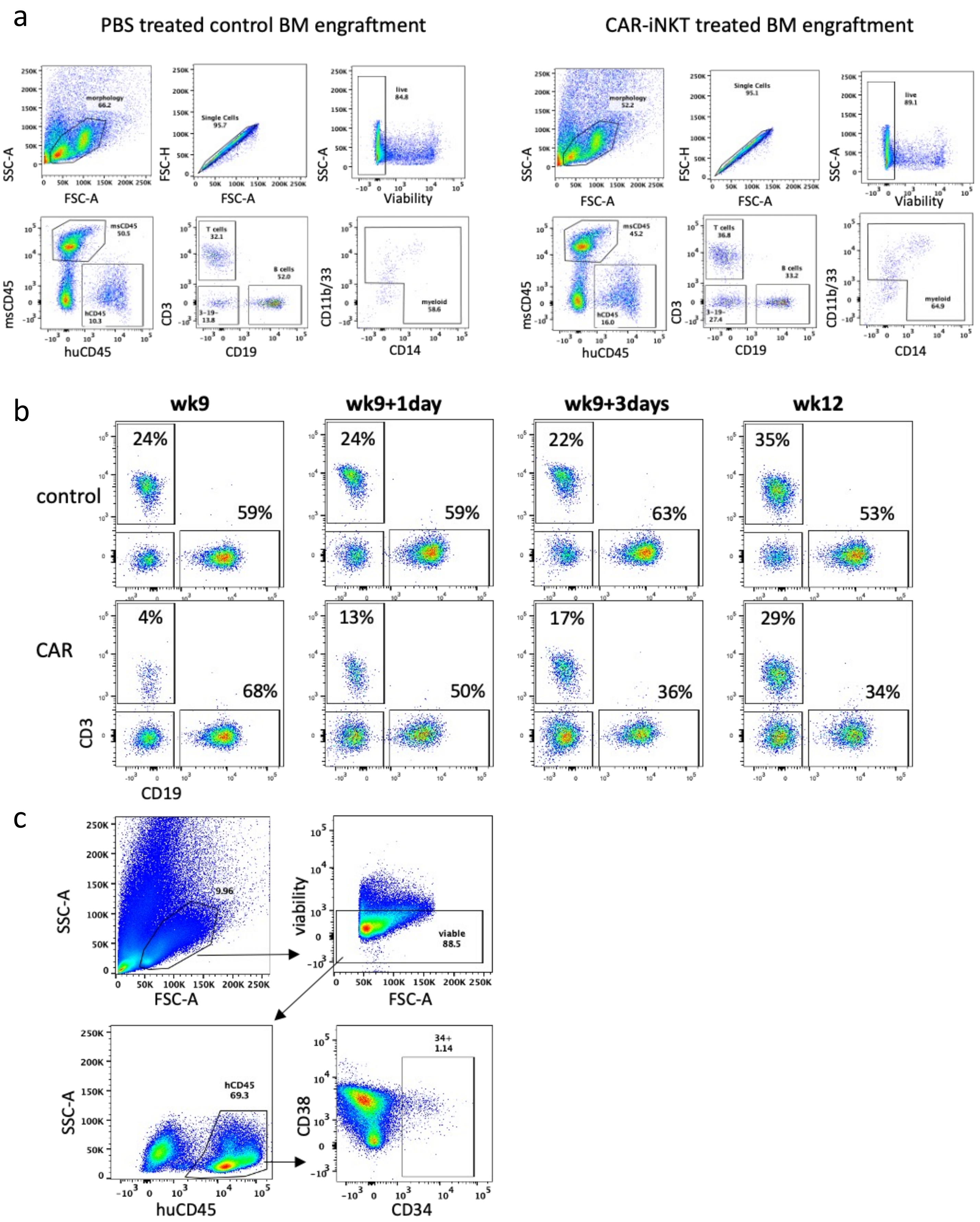


**Suppl Figure 10. Analysis of peripheral blood and bone marrow of humanized mice receiving bi-specific CAR-iNKT.** **a.** representative flow plots showing gating strategy used to determine peripheral blood (PB) and bone marrow (BM) engraftment and lineage output. The data shown is from BM at cull for PBS treated (left) and CAR-iNKT treated (right) mice. **b.** representative flow plots showing frequency of CD19+ B cells and CD3+ T cells in the PB of control mice treated with PBS (top row) and mice treated with CAR-iNKT at 9 weeks (bottom row). Data shown as % of hCD45+ cells. **c.** representative flow plots showing gating strategy used to determine immature CD34+ cells in the BM at cull. **d.** proportion of hCD45 cells in BM of secondary engrafted NSG mice. Each mouse was injected with 1M whole BM from primary engrafted mice treated with PBS (n=4) or 10M CAR (n=4).
